# Metagenomic methylation patterns resolve complex microbial genomes

**DOI:** 10.1101/2021.01.18.427177

**Authors:** Elizabeth G. Wilbanks, Hugo Doré, Meredith H. Ashby, Cheryl Heiner, Richard J. Roberts, Jonathan A. Eisen

## Abstract

The plasticity of bacterial and archaeal genomes makes examining their ecological and evolutionary dynamics both exciting and challenging. The same mechanisms that enable rapid genomic change and adaptation confound current approaches for recovering complete genomes from metagenomes. Here, we use strain-specific patterns of DNA methylation to resolve complex bacterial genomes from the long-read metagenome of a marine microbial consortia, the “pink berries” of the Sippewissett Marsh. Unique combinations of restriction-modification (RM) systems encoded by the bacteria produced distinctive methylation profiles that accurately binned and classified metagenomic sequences. We linked the methylation patterns of each metagenome-assembled genome with encoded DNA methyltransferases and discovered new restriction modification (RM) defense systems, including novel associations of RM systems with RNase toxins. Using this approach, we finished the largest and most complex circularized bacterial genome ever recovered from a metagenome (7.9 Mb with >600 IS elements), the finished genome of *Thiohalocapsa* sp. PB-PSB1 the dominant bacteria in the consortia. From these methylation-binned genomes, we identified instances of lateral gene transfer between sulfur-cycling symbionts (*Thiohalocapsa* sp. PB-PSB1 and *Desulfofustis* sp. PB-SRB1), phage infection, and strain-level structural variation.

## Introduction

In nature, bacterial and archaeal genomes are far from the tidy, static sequence of letters in our databases. They are, quite simply, alive – with all of the dynamism and complexity that we associate with life. Genomes can change substantially within the lifetime of a single cell, catalyzed by the intra- and inter-genomic shuffling of homologous recombination, mobile genetic elements, and phage. Unlike the gradual accumulation of point mutations, such bulk rearrangements can abruptly diversify an organism’s phenotypic traits and alter its niche (Bao et al 2016, Berube et al 2019, Doré et al 2020, Hehemann et al 2016, Rocap et al 2003). Horizontal gene transfer (HGT), such as the acquisition of pathogenicity islands or antibiotic resistance genes from a distantly related species, is perhaps the most notorious example of recombination abruptly changing an organism’s capabilities. However, even small-scale recombination within an organism’s own genome can alter important phenotypes, such as biofilm formation regulated by excision/insertion of an IS-element (Bartlett et al 1988, Higgins et al 2007, Ziebuhr et al 1999). The very same molecular features that enable rapid evolutionary change (genomic repeats, unusual sequence content and composition) also present analytical challenges, creating a disturbing blind spot in our study of microbial eco-evolutionary dynamics.

Metagenomic assembly algorithms founder when confronted with repetitive sequences; DNA sequences generated by most commonly used high-throughput methods are too short to unambiguously resolve the correct path through these complex regions of the assembly graph (Olson et al 2019). From samples with co-existing strains of the same species or for organisms primed for rearrangements because of their richness in repeats like transposons, we typically recover only genomic shrapnel, their recombination hotspots expunged. The highest quality metagenome assembled genomes (MAGs) often come from the most clonal species in a community (e.g. Banfield et al. 2017), not necessarily the most abundant or ecologically important (Chen et al. 2020). Assembly shortcomings beget further challenges, as smaller assembled sequences (contigs or scaffolds) are more difficult to correctly assign to their genomes of origin (i.e. binning).

Binning algorithms suffer from similar challenges because they classify assembled metagenomic sequences based on a set of shared, distinctive genome-wide signals (Chen et al 2020). Commonly used signals include phylogenetic profiles (sequence similarity to known organisms, e.g. Huson et al 2007), sequence composition (GC content or tetranucleotide frequency, e.g. Dick et al 2009, Tyson et al 2004), and relative abundance (coverage variation within a sample or across samples, e.g. Albertsen et al 2013). Accurate bins draw support from multiple, concordant signals that persist across all the sequences constituting the draft genome (Meyer et al 2018, Sieber et al 2018). However, in the mosaic genomes of many bacteria and archaea, such genome-wide consistency does not exist. Infecting (pro)phage, mobile elements, and laterally transferred genes all have evolutionary histories distinct from their host genome. The discord in phylogenetic and compositional profiles between these regions and the rest of the genome confounds binning algorithms relying on such signals (Maguire et al 2020). It remains challenging to faithfully reunite those sequence fragments that once comingled within the cell. Recent advances in binning leverage information about the genome orthogonal to its sequence, such as chromosomal conformation (Beitel et al 2014, Stewart et al 2018) or DNA methylation (Beaulaurier et al 2018).

In the present work, we studied the DNA methylation signals that bacteria and archaea use to discriminate their *own* genome from foreign DNA to overcome issues with assembling and binning complex microbial genomes from metagenomes. The most common base modifications in bacterial and archaeal genomes are made by the DNA methyltransferases (MTases), frequently associated with restriction-modification (RM) systems (Blow et al 2016). The restriction endonucleases of an RM system defend their host from foreign DNA by cleaving unmethylated DNA at sequence-specific recognition sites (Figure 1). The cognate MTase methylates recognition sites in the host’s genome, thereby protecting them from restriction enzyme activity (Figure 1C, Murray 2000). Beyond their role in host defense, MTases have been shown to play important physiological roles, from regulating gene expression to DNA replication and repair (Sanchez-Romero and Casadesus 2020).

**Figure 1.**
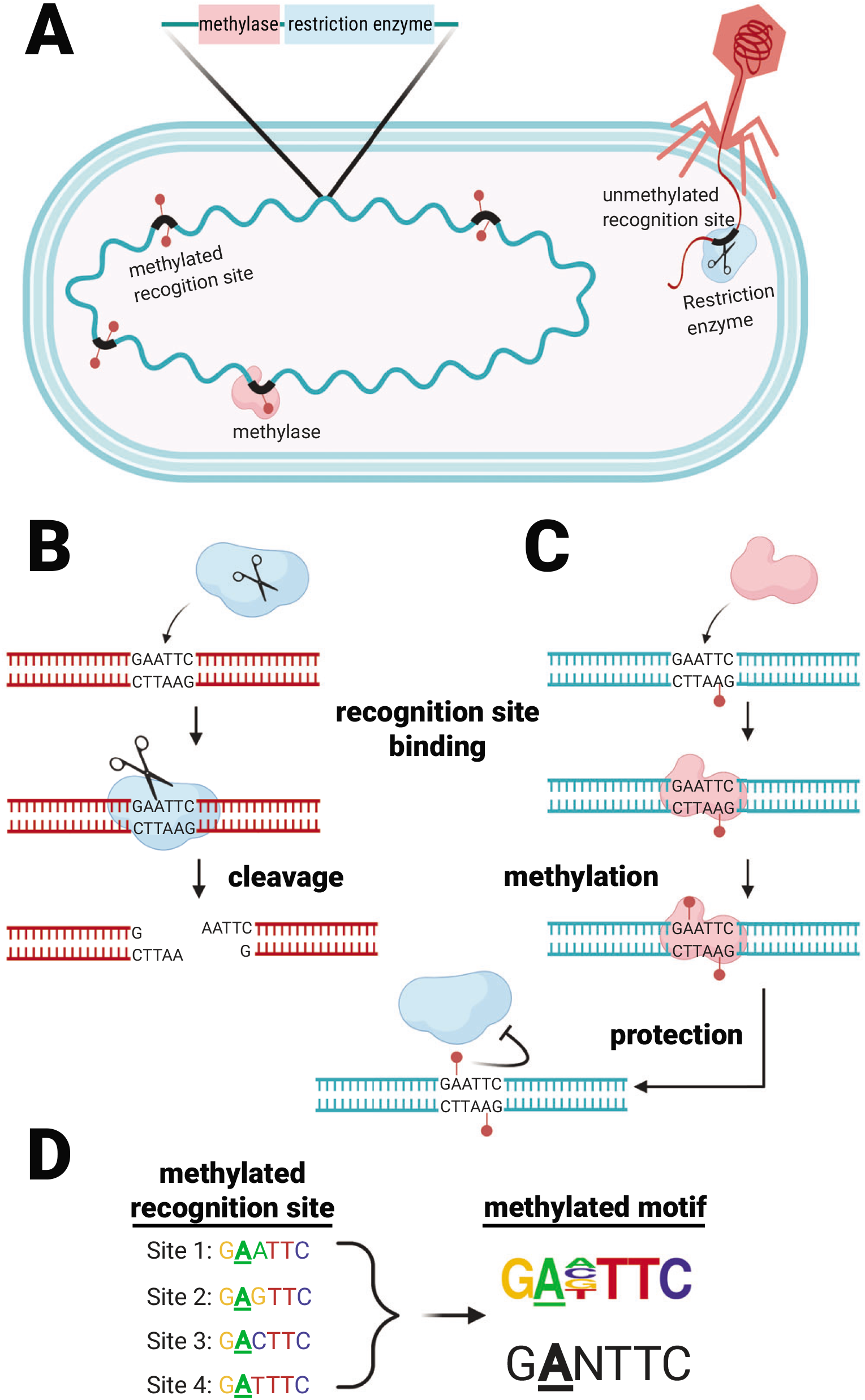
Restriction-modification (RM) systems provide bacteria and archaea with a defense against foreign DNA by discriminating self from non-self DNA based on methylation patterns. **(A)** RM systems, such as the Type II RM illustrated here, consist of a methylase (pink) and restriction enzyme (blue) which both recognize short, specific sequences in the genome (“recognition binding sites”, thick black lines). **(B)** Unmethylated recognition sites, such as the example shown in an infecting phage genome, are cleaved by the restriction enzyme. **(C)** The methylase acts like an antitoxin, an antidote to the restriction enzyme’s toxicity. The methylase binds and methylates the recognition site (which is often palindromic) at a specific base. The methylase binds and modifies hemi-methylated recognition sites, where one but not both strands are unmethylated, a characteristic which helps the cell discriminate newly replicated host DNA (hemi-methylated) from completely unmethylated foreign DNA. Methylation of both 5’ and 3’ strands in the recognition site inhibits the cognate restriction enzyme, protecting that site from cleavage. **(D)** Methylases (and their cognate restriction enzymes) often tolerate variation in some positions of their recognition sites, as shown for position 3, in this example. A methylase’s binding site sequence can be discovered by analyzing the sequence context around methylated bases in the genome, and summarized by a sequence motif where the methylated base is underlined (shown here as a sequence frequency logo, top, or a consensus sequence, bottom).

Specific MTase recognition sites can be discovered from genome-wide surveys of DNA modification by examining the short stretches of sequence surrounding the methylated base, and summarizing recurrent patterns as methylated motifs (Figure 1D, Beaulaurier et al 2019). RM systems are diverse and widespread amongst bacteria and archaea, and like many defense systems, they vary greatly even between closely related species (Koonin et al 2017). We identified species-specific methylation patterns on metagenomic contigs from the DNA polymerase kinetics of Pacific Biosciences (PacBio) sequence data. We used this methylation information to bin and assemble complex bacterial genomes from a microbial consortium, the “pink berries” of the Sippewissett marsh, macroscopic microbial aggregates, for which we previously recovered complete but highly fragmented MAGs (Wilbanks et al 2014).

## Results

### Methylation in metagenomes: detection and clustering of sequence data

PacBio data from “pink berry” aggregates were assembled to produce 18 megabases (Mb) of sequence on 169 contigs, with an N50 of 413 kb, where the largest contig was 3.5 Mb in size. This assembly recruited back 87% of the error corrected reads, indicating that it was a reasonable representation of the data. N^6^-methyladenine (6mA) was detected on every contig (modification QV ≥□20, *i.e*. p-value ≤ 0.01), while N^4^-methylcytosine (4mC) was detected on 152 out of the 169 contigs (Supplemental Figure 1 and Supplemental Data 1). The average frequency of m6A detections per 10 kb of contig sequence was independent of coverage above 40x coverage, indicating good detection sensitivity for most assembled contigs (Supplemental Figure 2A). In contrast, m4C modifications were both rarer and strongly correlated with coverage up to ~60x coverage, which suggests decreased detection sensitivity on many contigs (Supplemental Figure 2B).

Thirty-two sequence motifs were identified from the sequence context of these methylations by analyzing a subset of large contigs in the dataset using the SMRT Analysis workflow (Supplemental Table 1). For each of these motifs, we quantified how many times the sequence occurred on a contig and whether that sequence was methylated. Thus, each contig has a “methylation profile” composed of 32 distinct methylation metrics, quantifying proportion of a motif’s occurrences that were methylated.

### Methylation-based clustering recovers metagenome-assembled genomes

The methylation profiles differed significantly between metagenomic contigs, and hierarchical clustering partitioned this data into seven distinct groups (Figure 2). These groups were largely recapitulated by independent clustering using t-distributed stochastic neighbor embedding (t-SNE) of the methylation profiles, and these groups represented taxonomically coherent bins of the dominant organisms in the consortia (Figure 3A, Table 1). These methylation clusters were also largely consistent with similarities in sequence composition, such as tetranucleotide frequency and GC-content (Figure 3B and Supplemental Figure 3, respectively).

**Figure 2.**
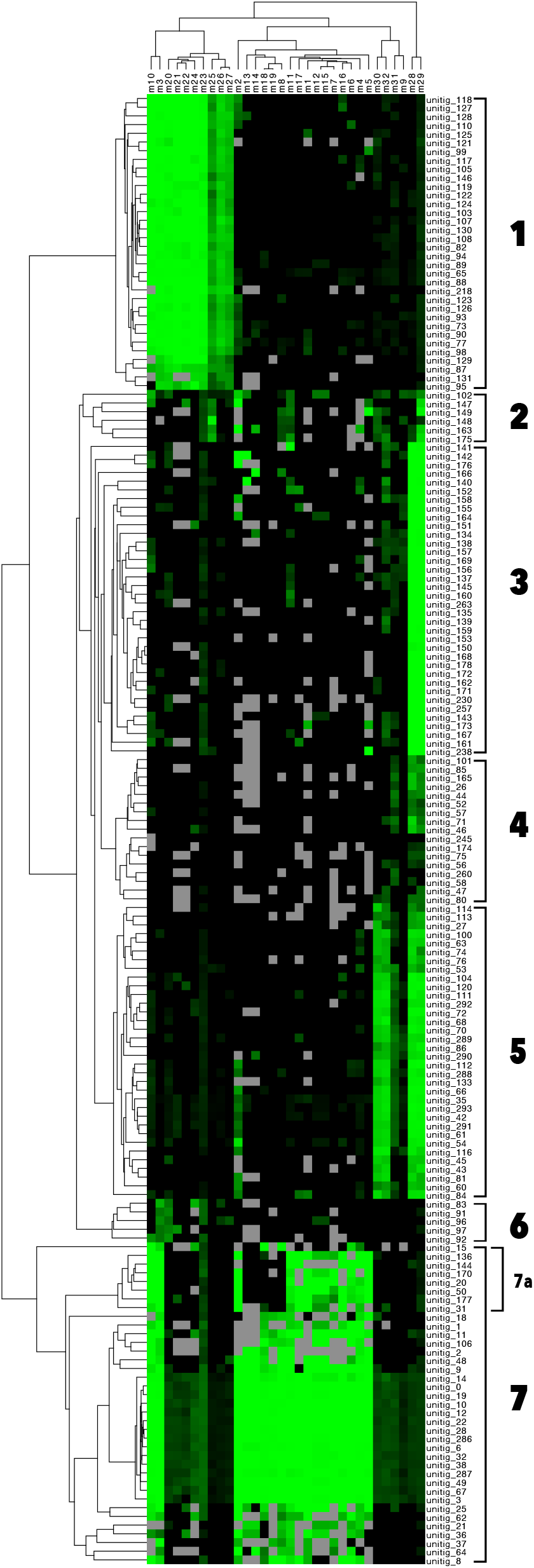
Patterns of DNA methylation group metagenomic contigs into distinct clusters via hierarchical clustering. The methylation status of 32 distinct sequence motifs (m1-m32, columns) is shown on every metagenomic contig (rows, unitig 1 - unitig 169). The value plotted is the percentage of motifs methylated on every contig (square root transformed); bright green color indicates a motif for which every instance on that contig was methylated (100%), and black shows motifs for which no instances were methylated on that contig (0%). When no instances of the sequence motif were observed on a contig, this is indicated as missing data (gray). Rows and columns have been hierarchically clustered based on Euclidean distance. Distinct methylation clusters have been numbered 1-7.

**Figure 3.**
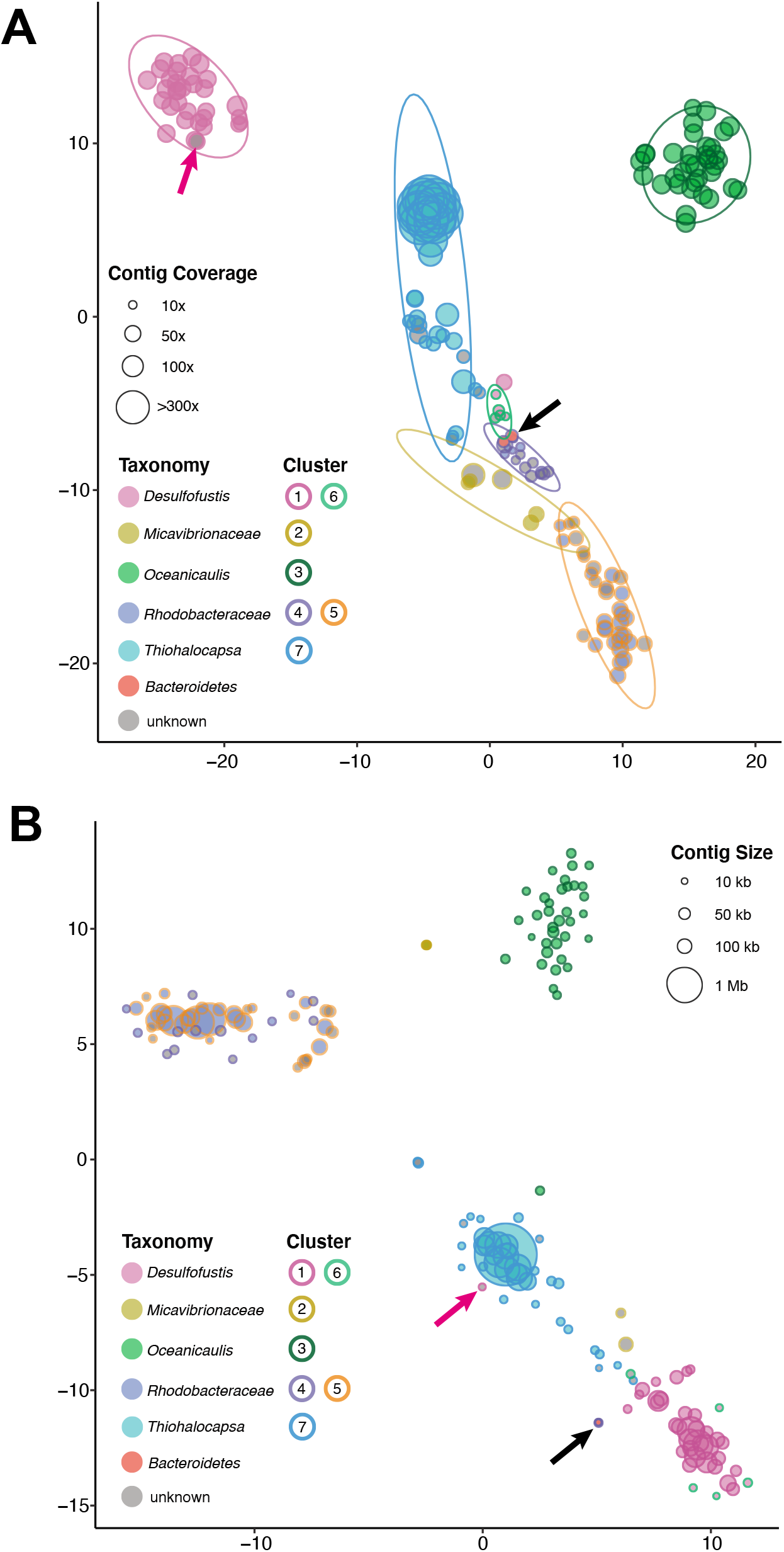
Visualization with t-distributed stochastic neighbor embedding (t-SNE) of **(A)** methylation profiles and **(B)** tetranucleotide frequencies create taxonomically distinct clusters. Point size is scaled to either contig coverage **(A)** or contig length **(B)**, fill color corresponds to the taxonomic assignment, and outline color represents the methylation-based hierarchical clusters defined in Figure 2. Prediction ellipses in panel A were defined for hierarchical methylation clusters with the assumption that the population is a multivariate t-distribution. Black arrows indicate the three overlapping low coverage, low GC (<45%) contigs within methylation cluster 4 that represent contamination from the *Bacteroidetes*. Pink arrows indicate a contig which had discordant binning between methylation profiling and tetranucleotide frequency analyses.

**Table 1.**
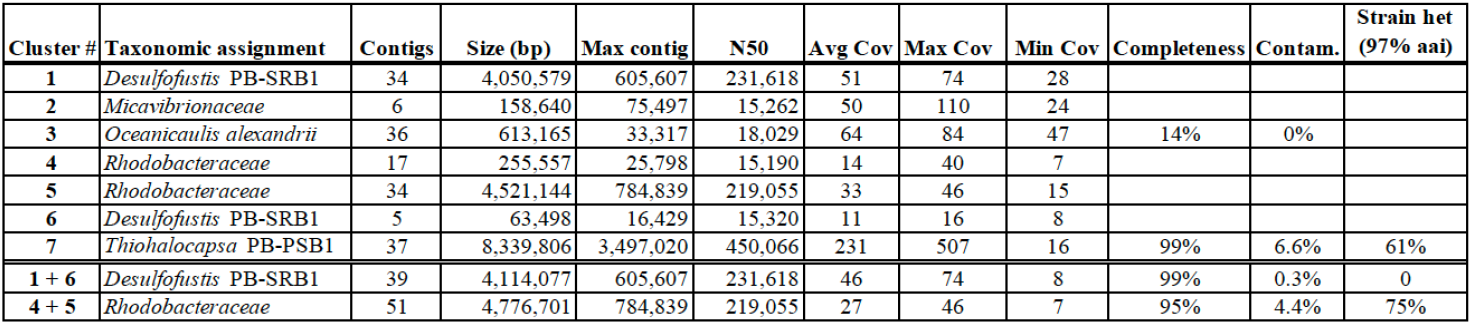
Summary of the seven methylation-based hierarchical clusters of metagenomic contigs defined in Figure 2. Metagenome-assembled genomes (MAGs) with completeness >90% are represented by methylation cluster 7, clusters 1+6, and clusters 4+5. Completeness was assessed by presence of lineage-specific single copy marker genes, while contamination was assessed by their presence in >1 copy. The proportion of observed multicopy marker gene sets sharing ≥97% amino acid identity is represented in the “strain heterogeneity” metric. For example, 16 marker genes in the *Rhodobacteraceae* MAG (clusters 4+5, bottom line in table) were found in duplicate copies. In 12 of these 16 genes though, their duplicate copies shared ≥ 97% aa identity, indicating these “contaminants” derived from highly similar strains or incomplete assemblies, rather than inclusion of distant organisms due to binning errors (e.g. 75% strain het. = 12/16).

### Binning and circular assembly of the Thiohalocapsa sp. PB-PSB1 genome (Cluster 7)

Cluster 7, the largest of the methylation groups at 8.3 Mb, represented a 99% complete MAG for *Thiohalocapsa* sp. PB-PSB1, the most abundant organism in the consortia (Table 1) (Wilbanks et al., 2014, Seitz et al. 1993). This bin assembled to long contigs (N50 450 kb, max 3.5 Mb), unlike corresponding Illumina MAGs, which were far more fragmented (N50 ~40 kb, max 160 kb). Contigs in this cluster larger than 100 kb (n=15) had an average coverage of 489x, while the smaller contigs (<25 kb, n=22) were lower coverage with an average of 57x.

Of the 37 contigs in this bin, 31 were clearly identified by sequence similarity as *Thiohalocapsa* sp. PB-PSB1 (Figure 3A). Six contigs did not have clear taxonomic assignments, but grouped most closely with other PB-PSB1 contigs based on sequence composition (Figure 3B). Five of these taxonomically unidentified contigs also shared strong assembly graph connectivity with other PB-PSB1 contigs. Contamination for this cluster, estimated based on the percentage of single copy marker gene sets present in multicopy, was predicted to be 6.6%. However, 61% of these multicopy marker genes (20 in total) shared ≥97% amino acid identity between copies, and no multicopy genes revealed hits to distantly related taxa. These multicopy genes, therefore, may not indicate contamination, but rather strain-level variation, incomplete assembly, or recent duplications.

Within methylation cluster 7, a subclade of seven, smaller contigs (7a, length < 25 kb, coverage 46 + 7x) shared an unusual methylation profile relative to other contigs. While the contigs in clade 7a encode the sequence of the characteristic PB-PSB1 motifs m8, m13, m14, m18, and m19, these motifs were rarely methylated. By contrast, these motifs were almost universally methylated when they occurred on the other contigs in methylation group 7, even on contigs with less than 40x coverage.

Reassembly of cluster 7 sequence data produced a circular assembly graph formed by 9 backbone contigs. In addition, there were 51 small contigs forming “bubbles” or spurs connected to this main assembly graph (length <25 kb, 543 kb total), and four “singleton” contigs unconnected to the circular assembly graph (length < 22 kb, 57 kb total). With manual curation, the genome was closed to produce a single circular contig of 7.95 Mb which represents the finished genome of *Thiohalocapsa* sp. PB-PSB1 (CP050890). This finished genome contains 606 insertion sequence (IS) elements which comprise 9.4% of the total genome sequence (Table 2). These IS elements were both diverse, belonging to 17 phylogenetically distinct families, and highly repetitive as demonstrated by a 1.5 kb IS154 transposon found in 44 identical copies distributed throughout the genome.

**Table 2.** A total of 606 insertion-sequence (IS) elements were identified in the finished, circularized genome of *Thiohalocapsa* sp. PB-PSB1. IS elements were classified into phylogenetically distinct families (based on the ACLAME database), and the total instances of each class was enumerated. The total number of nucleotides within each IS element class was determined and its percentage of the complete genome computed.

| family | # | nucleotides | % of genome |
| --- | --- | --- | --- |
| IS4 | 190 | 174,828 | 2.2 |
| IS91 | 120 | 172,158 | 2.17 |
| ISL3 | 54 | 65,064 | 0.82 |
| IS5 | 50 | 52,601 | 0.66 |
| ISAS1 | 35 | 40,445 | 0.51 |
| IS630 | 29 | 36,442 | 0.46 |
| ISAZO13 | 18 | 26,086 | 0.33 |
| IS110 | 16 | 27,959 | 0.35 |
| IS66 | 16 | 32,674 | 0.41 |
| IS1634 | 15 | 28,613 | 0.36 |
| IS21 | 15 | 30,347 | 0.38 |
| IS200/IS605 | 14 | 8,291 | 0.1 |
| IS1182 | 10 | 16,013 | 0.2 |
| ISKRA4 | 9 | 15,814 | 0.2 |
| IS701 | 7 | 12,957 | 0.16 |
| IS3 | 4 | 4,224 | 0.05 |
| ISNCY | 3 | 2,815 | 0.04 |
| new | 1 | 1,846 | 0.02 |
| **total** | **606** | **749,177** | **9.42** |

### Identifying HGT in Desulfofustis sp. PB-SRB1 (Clusters 1 & 6)

Methylation clusters 1 and 6 comprised the complete (99%) and uncontaminated (<0.5%) genome of *Desulfofustis* sp. PB-SRB1. Sequence similarity and composition both confirmed that these two methylation clusters contain contigs originating from a single organism that is highly similar to prior data from *Desulfofustis* sp. PB-SRB1 (Figure 3B) (Wilbanks et al 2014). Cluster 1 shared some common methylated motifs with PB-PSB1 (e.g. m3 and m10), but also contains other frequently methylated motifs (m20-m27) that were unique to PB-SRB1 (Figure 2). Though the methylation profiles of contigs in clusters 1 and 6 differ from one another (Figure 3A), they shared several key similarities, namely, the methylation of motifs m20, m3 and m21 and absence of methylation on other motifs (Figure 2). These two clusters differed significantly in coverage: cluster 1 contained the majority of the genome at ~ 50x coverage, while cluster 6 contained only 63 kb at ~11x coverage (Table 1). Many cluster 6 contigs shared sequence similarity with larger higher coverage portions of the assembly grouped in cluster 1, and may represent structural variants (some had transposon deletions or sequence rearrangements relative to their parent contigs).

One 10 kb contig, unitig_146, grouped with cluster 1 in both hierarchical and t-sne clustering based on methylation profiling, but was most similar to *Thiohalocapsa* sp. PB-PSB1 contigs in sequence composition (Figure 3). Given the conflicting evidence, we further investigated this contig to determine whether this represented HGT or a binning error. We manually inspected this contig in the reassembled PB-SRB1 genome and found no evidence of misassemblies. This contig encodes two class C beta-lactamase genes alongside a D-glutamate deacylase and prolidase, functions which suggest that this gene cassette enables both the opening of beta-lactam rings and decarboxylatation to their constituent D-amino acids (Figure 4). Flanking these genes were two transposons (IS481 and IS701) most closely related to homologs from the *Desulfobacterales* and found in multiple copies on the other contigs in the PB-SRB1 genome. Neither of these transposons were found in the closed genome of *Thiohalocapsa* sp. PB-PSB1. This contig was in a complex region of the PB-SRB1 assembly graph, with connectivity to two large contigs containing PB-SRB1 marker genes (>100 kb), and two smaller contigs (<10 kb). Alignment of these contigs and their component reads revealed numerous structural variants in this region (duplications and inversions). Combined with the distinctive methylated motifs present on this contig, these findings give us confidence in our assignment of this contig as a true portion of the PB-SRB1 genome.

**Figure 4.**
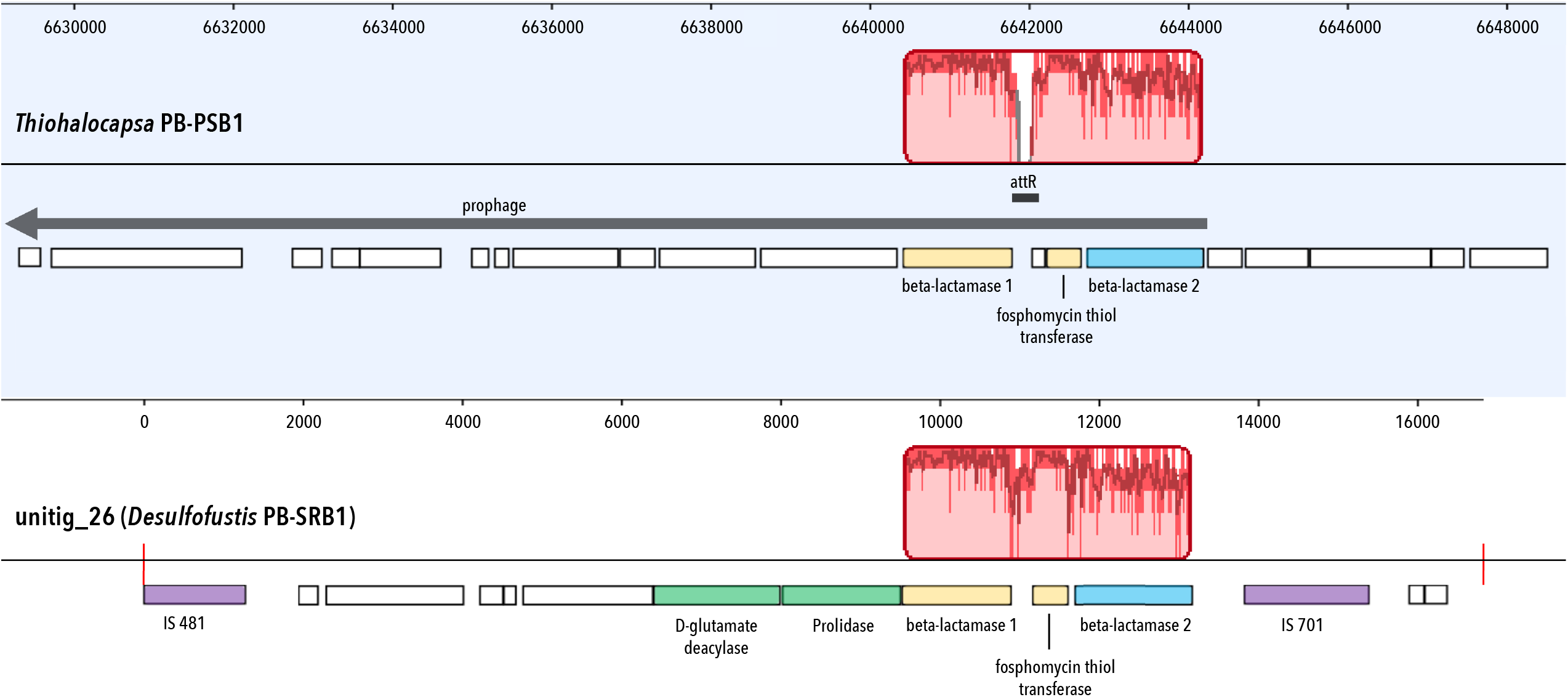
Genome alignment shows evidence for the horizontal transfer of antibiotic resistance genes between the bacterial symbionts *Thiohalocapsa* sp. PB-PSB1 (top, finished genome) and *Desulfofustis* sp. PB-SRB1 (bottom, unitig_26). Highlighted in red is the homologous region identified by whole genome alignment, where bar height represents the degree of conservation. Highlighted in yellow are the highly conserved genes: beta-lactamase 1 (88% nucleotide identity; 97% aa similarity) and a fosphomycin resistance thiol transferase (91% nt id; 97% aa similarity). Beta-lactamase 2 (in blue), which contained an N-terminal twin arginine leader peptide, was less closely related (74% nt id; 88% aa sim). On unitig_26, this region was flanked by transposons (purple) found on several other contigs in the *Desulfofustis* PB-SRB1 assembly. In the *Thiohalocapsa* sp. PB-PSB1 genome, this region falls within a 29 kb prophage (grey arrow). The attR insertion site (black line) for the prophage is not conserved in the unitig_26 sequence, as evidenced by the dip in sequence similarity in this region.

Alignment of this *Desulfofustis s*p. PB-SRB1 contig with the closed *Thiohalocapsa* sp. PB-PSB1 genome revealed sequence similarity only in the 3.7 kb region containing these two beta-lactamase genes (Figure 4). This region of the PSB1 genome overlaps with a 29 kb prophage complete with flanking attL and attR insertion sites. However, this prophage region was not conserved in PB-SRB1.

### *Resolving three distinct* Alphaproteobacteria

The remaining methylation clusters 2 – 5 are composed of contigs from 3 different *Alphaproteobacteria*. Motif m29 (G<u>A</u>NTC) was frequently methylated on nearly all contigs from clusters 2 – 5. Cluster 3, which was characterized by frequent methylation of motif m28 (RG<u>A</u>TCY) in addition to m29, represents a partial and uncontaminated MAG closely related to *Oceanicaulis alexandrii* (Figure 2–3, Table 1).

### *A novel* genus in the Rhodobacteraceae (*Clusters 4&5*)

Binning together methylation clusters 4 and 5, we recovered a 4.7 Mb, 95% complete MAG with 4% contamination corresponding to strain heterogeneity (Table 1). This long-read bin shared 99.8% ANI with a 4.2 Mb MAG from our Illumina dataset (PB-A2), estimated to be 97.5% complete with 0.4% contamination. By tree placement and ANI (99%), these MAGs’ closest relative in public databases is UBA10424 (GCA_003500165.1, N50 = 13 kb), an 88% complete MAG extracted from our previous, lower coverage sequencing of this same system in 2010, and proposed to be the sole representative of a novel genus in the *Rhodobacteraceae*.

While these clusters were separated in methylation space, they grouped closely together based on sequence composition (Figure 3). Cluster 5 contained the majority of the genome (4.5 Mb), while cluster 4 contained smaller, lower coverage fragments (Table 1). Many of the high GC contigs in these methylation clusters could be identified by sequence similarity as belonging to the family *Rhodobacteraceae*. Contigs without clear taxonomic identity could be linked with the other *Rhodobacteraceae* contigs based on their overlap-based assembly graph connectivity. These groups shared methylation of m28 and m29 with cluster 3 (*Oceanicaulis*) but were distinguished by the methylation of m30 (CANC<u>A</u>TC) and m32 (GATGG<u>A</u>).

Cluster 4 contained three low GC contigs from the *Bacteroidetes* that represent contamination (black arrow Figure 3). Though these contigs did contain detected modifications (Supplemental Data 1), these methylations either never (unitig_245) or rarely (unitig_260, unitig_174) occurred within one of the 32 characteristic motifs. This data suggests that these contigs grouped with the lowest coverage contigs (cluster 4) in our dataset based on the absence of methylated motifs, rather than any positive signal.

### *Linking phage infection with a novel* Micavibrionaceae *species (Cluster 2)*

Cluster 2 comprises 158 kb of sequence on six contigs, four of which were identified as belonging to the *Micavibrionaceae* by sequence similarity (Table 1). The methylation profile of cluster 2 contained m29, like the other *Alphaproteobacteria*, but was missing m28. Cluster 2 was further distinguished by distinctive combination of methylated motifs m25, a 4mC motif (CCAGCG), and m11 (GAGATG). The contigs identified as*Micavibrionaceae* (30x coverage) mapped with high identity to a MAG (PB-A3) binned from our parallel Illumina assembly (84% complete, 0.5% contamination, N50 32 kb). This MAG’s closest relative in public databases is UBA10425 (GCA_003499545.1), an 80% complete genome extracted from our prior, lower coverage sequencing of this same system, and proposed to be the sole representative of a novel genus within the *Micavibrionaceae*.

The remaining two contigs in cluster 2 were present at significantly higher coverage (70x and 110x) and were identified as putative phage sequences. Notably, while these contigs clustered closely with the others based on their methylation profiles (Figure 3A), they had markedly different sequence composition relative to the other *Micavibrionaceae* contigs (Figure 3B). The first of these, unitig_102, was an outlier at 110x coverage, which was the highest coverage contig in this dataset that was not from *Thiohalocapsa* sp. PB-PSB1. This 75 kb contig is predicted to encode a complete *Siphoviridae* dsDNA phage genome, with both structural and DNA replication genes. Ten of these coding sequences shared high percent identity (32-66% aa identity) with a cultured temperate phage, phiJI001, known to infect an alphaproteobacterial isolate from the genus *Labrenzia*. Searches of this *Siphoviridae* contig against our Illumina based MAGs found high percentage identity matches to several contigs binned to *Micavibrionaceae* PB-A3. These contigs were linked to the PB-A3 bin based on paired-end read connectivity, but not by our sequence composition or coverage-based analyses. Unitig_102 could be circularized (with manual trimming), a common characteristic of *Siphoviridae* genomes; however, the PacBio data and Illumina paired end reads both supported scaffolding with a 100kb contig in the Illumina assembly which contained *Micavibrionaceae* marker genes.

### Novel restriction-modification (RM) systems and orphan methyltransferases explain the diversity of methylation patterns in the metagenome

To further investigate the patterns of DNA methylation in the consortia, we analyzed each MAG individually which detects methylated motifs with greater sensitivity. For the incomplete genomes (e.g. the alphaproteobacteria), we also analyzed their corresponding Illumina-assembled MAGs as validation. The genomes each contained from 4 to 17 different methylated motifs, and every genome had at least one methylated motif unique to that organism in the consortia (Figure 5A). This analysis recovered 30 of the 32 motifs from our initial prediction, and also discovered 13 additional methylated motifs (Supplemental Data 2). There is substantial novelty in these genome modifications: 40% of these methylated motifs (n=17; red and navy bars in 4C) have never been reported in genome-wide methylation studies or databases of RM recognition sites (Roberts et al 2015).

**Figure 5.**
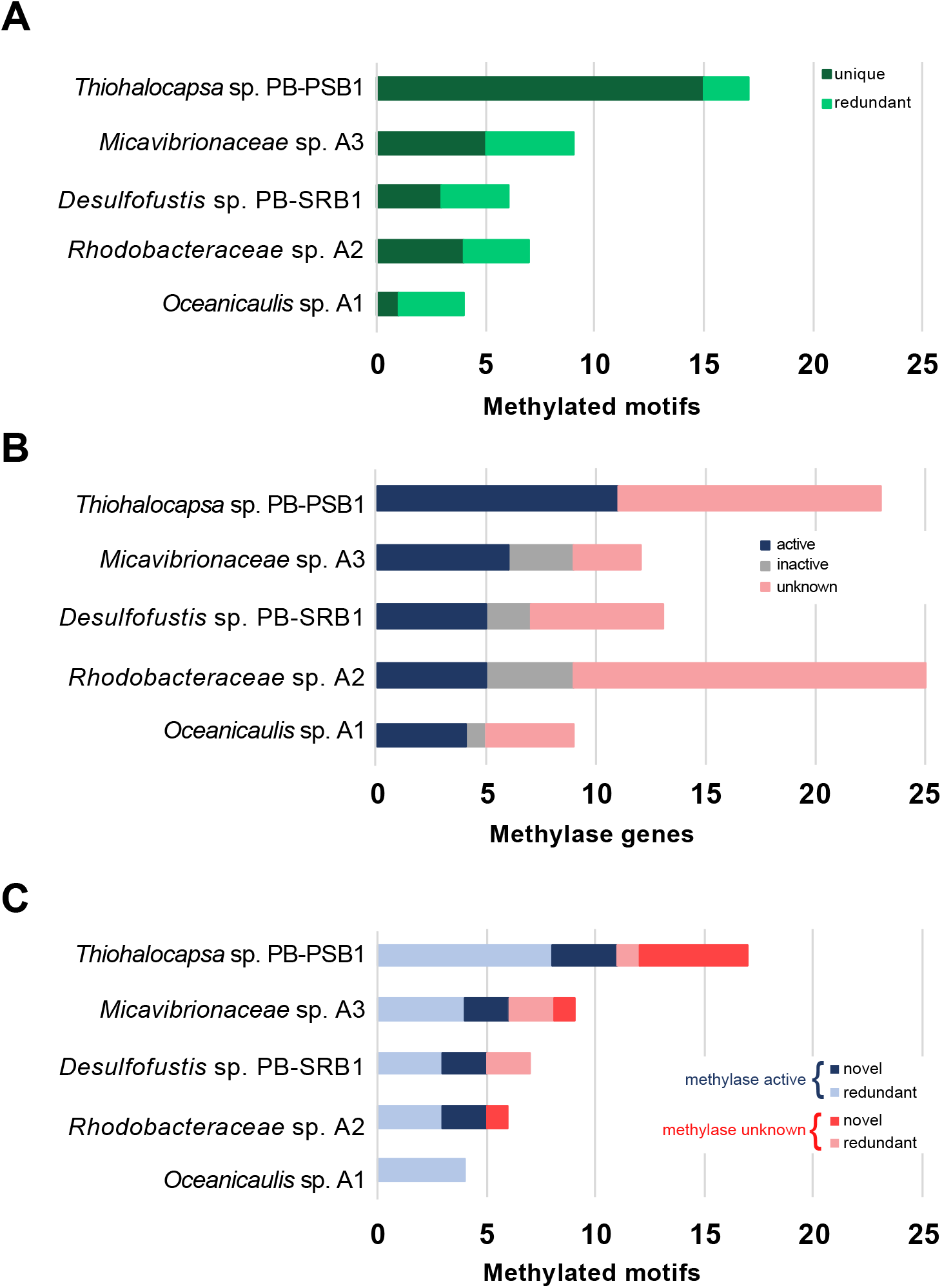
**(A)** Analysis of each metagenome-assembled genome (MAG) demonstrates that while some methylated motifs were observed amongst several consortia members (*redundant*, light green), each MAG contained methylated motifs unique to that species in the dataset (*unique*, dark green). **(B)** Each organism contained numerous methylase genes, which were classified as “active” (navy blue bars) when methylase’s predicted recognition sequence was methylated, “inactive” (grey) where the predicted recognition sequence was not frequently methylated, or “unknown” (pink) if the recognition sequence could not be predicted bioinformatically. **(C)** Most methylated motifs in each MAG could be linked with a predicted source methylase (light blue, navy), though each genome (except for *Oceanicaulis*) had some motifs for which the source methylase remains unknown (pink, red). All genomes except for *Oceanicaulis* sp. A1 contained novel motifs yet to be documented in REBASE (red, navy), while others were redundant with known RM recognition sequences (light blue, pink).

We investigated the source of these methylation patterns by annotating the MTase and restriction enzyme genes in each genome. We found between 9 and 24 different MTase genes in each genome, and for ~50% of these genes, we could bioinformatically predict their recognition sequences, many of which matched methylated motifs in the genomes (Figure 5B, Supplemental Data 2). Every genome, except for *Oceanicaulis*, encoded 2-3 novel RM systems which we predict recognize and methylate (or cut) novel sequence motifs (navy blue bars, Figure 5C).

Several MTases in the *Oceanicaulis* and *Rhodobacteraceae* MAGs were found to be encoded on putative phage or prophage contigs (3 and 8 MTases, respectively). Most of these phage MTases occurred as “orphans”, without a corresponding restriction enzyme, though one 60kb phage contig in the *Rhodobacteraceae* MAG encoded 3 Type II orphan MTases, as well as a complete Type I RM system (Supplemental Data 2). These sequences were quite divergent from known MTases, and as such their recognition sites could rarely be predicted, with the exception of G<u>A</u>TC phage MTases which would likely confer protection against the hosts’ RGATCY cleaving restriction enzymes.

Examining the RM systems in the *Thiohalocapsa* PB-PSB1 genome, we discovered that RM genes frequently co-occurred with putative RNase or RNA interferase toxin genes from the *vapC* or *hicA* family. Six of the 23 MTases in this genome were immediately flanked by these *vapC* or *hicA* toxins. Five of these cases encoded complete RM systems – including 3 out of the 4 complete Type I operons in the genome (Supplemental Data 2). These loci encoded only the toxin gene without an antitoxin; however, *vapB* and *hicB* family antitoxins were found elsewhere in the genome.

## Discussion

Examining metagenomic methylation patterns, we binned and assembled complex bacterial genomes from a microbial consortium with substantial strain variation. Such methylation-based binning has been tested using cultured mock communities and the mouse gut microbiome (Beaulaurier et al 2018); however, this approach has yet to be validated in other systems. Though we use a different workflow in identifying methylated motifs, we similarly found that methylation patterns faithfully distinguish contigs from distinct species. We identified the host for a complete phage genome based on their similar patterns of DNA methylation, the first application of this novel approach for linking phage with their hosts.

With our approach, we finished the largest and most complex circular bacterial genome yet recovered from a metagenome. Though closing genomes is now routine with bacterial and archaeal isolates, circularized metagenome assembled genomes (cMAGs) remain rare and tend to be both clonal and small (Chen et al 2020), though they are becoming increasingly accessible with long read sequencing (Moss et al 2020). At 7.9 Mb, the circularized genome of *Thiohalocapsa* PB-PSB1 is the largest finished genome ever reconstructed from a metagenomic sample, exceeding long-read pseudomonad cMAG by nearly 1.5 Mb (White et al 2016).

Previous short read metagenomes recovered complete but highly fragmented genomes for the most abundant species in the consortium, *Thiohalocapsa* sp. PB-PSB1 and *Desulfofustis* sp. PB-SRB1 (Wilbanks et al 2014), which suggested strain complexity or intragenomic repeats. Indeed, the finished *Thiohalocapsa* PB-PSB1 genome is highly repetitive: it harbors amongst the highest number of transposons ever reported in a bacterial or archaeal genome (Newton and Bordenstein 2011, Touchon and Rocha 2007). With 9.4% of its genome comprising transposon sequence, *Thiohalocapsa* PB-PSB1 has an unusual genome structure for free-living bacteria, though not unprecedented among aggregate- and bloom-forming phototrophs (Hewson et al 2009, Kaneko et al 2007). Repetitive mobile elements are not only vehicles for transposition and HGT, but also frequently flank hotspots of homologous recombination in bacterial genomes (Everitt et al 2014, Oliveira et al 2017). The transposon abundance in *Thiohalocapsa* PB-PSB1, thus, indicates substantial potential for recombination and genome plasticity.

Strain-level structural variants of transposons (e.g. deletions, inversions) were evident in both the PB-PSB1 assembly graph and in mapped reads spanning transposon regions in the finished genome. In the hierarchical clustering of contigs by methylation profile, we observed a clade of small contigs from PB-PSB1 where several distinctive sequence motifs remained unmethylated (Figure 2, cluster 7a). These sequences were structural variants of the finished, circular genome and contained transposons which we found, in different sequence contexts, elsewhere in the finished genome. Considered together, this evidence suggests that sequences in cluster 7a originate from a distinct strain, distinguished from the most abundant PB-PSB1 strain by genome rearrangements near transposons, and missing or inactive MTases. While these missing methylations could be an artifact of low coverage, we find this interpretation unlikely as these motifs were frequently methylated on many lower coverage contigs in PB-PSB1 and coverage as low as 15x can reliably detect Type I motif methylations (Blow et al. 2016).

The *Desulfofustis* sp. PB-SRB1 genome was complete but remained draft quality (n=72, N50 385 kb, max 930 kb), due to strain-level structural variants and lower coverage. Methylation profiling provided key information allowing us to link an island of horizontally transferred antibiotic resistance genes to the *Desulfofustis* sp. PB-SRB1 genome. This small contig would have almost certainly been erroneously binned with the *Thiohalocapsa* sp. PB-PSB1 genome by most algorithms; however, we were able to correctly identify it as belonging to *Desulfofustis* sp. PB-SRB1 based on its distinctive methylation profile.

The patterns of methylation in the pink berry MAGs are highly novel and offer a window into unexplored microbial DNA methylation systems: 40% of methylated motif we found have no matches in restriction enzyme databases (Roberts et al. 2015). Systematically annotating the MTase genes in each genome, we discovered 7 RM systems that we predict recognize some of these novel methylated motifs. The majority of these novel MTases were Type I systems recognizing asymmetric target sites with the nonspecific spacer of 4 - 8 bp (typically 6 bp), characteristic of Type I RM systems (Murray 2000). *Thiohalocapsa* sp. PB-PSB1 remains a rich target the discovery of yet more novel MTases, with a dozen uncharacterized MTases (pink bars in Figure 5B) and five novel methylated motifs without a predicted MTase (red bars in Figure 5C). Clearly, further experimental characterization of these MTases and restriction enzymes is warranted and could yield enzymes of biotechnological utility (Buryanov and Shevchuk 2005).

In the PB-PSB1 genome, we discovered RNA-targeting *vapC* and *hicA* toxin genes immediately adjacent to RM systems, a co-occurrence that has not previously been reported. We propose that these VapC and HicA homologs play a role in programmed cell death, analogous to the PrrC-Ecoppr1 abortive infection system in *E. coli* (Tyndall et al 1994). Though these systems do not show homology based on sequence comparisons, the functional parallels are notable. The PrrC abortive infection system includes an anticodon tRNA nuclease which initiates programmed cell death, should the Type I restriction enzyme defense fail against phage infection (Figure 6) (reviewed by Kaufmann 2000). Our preliminary investigations found *vapC* or *hicA* homologs also co-occurred with RM genes in other bacterial genomes. Such RNA-acting apoptotic toxins may be more widely integrated with restriction enzymes as a “fail-safe” defense than was previously appreciated.

**Figure 6.**
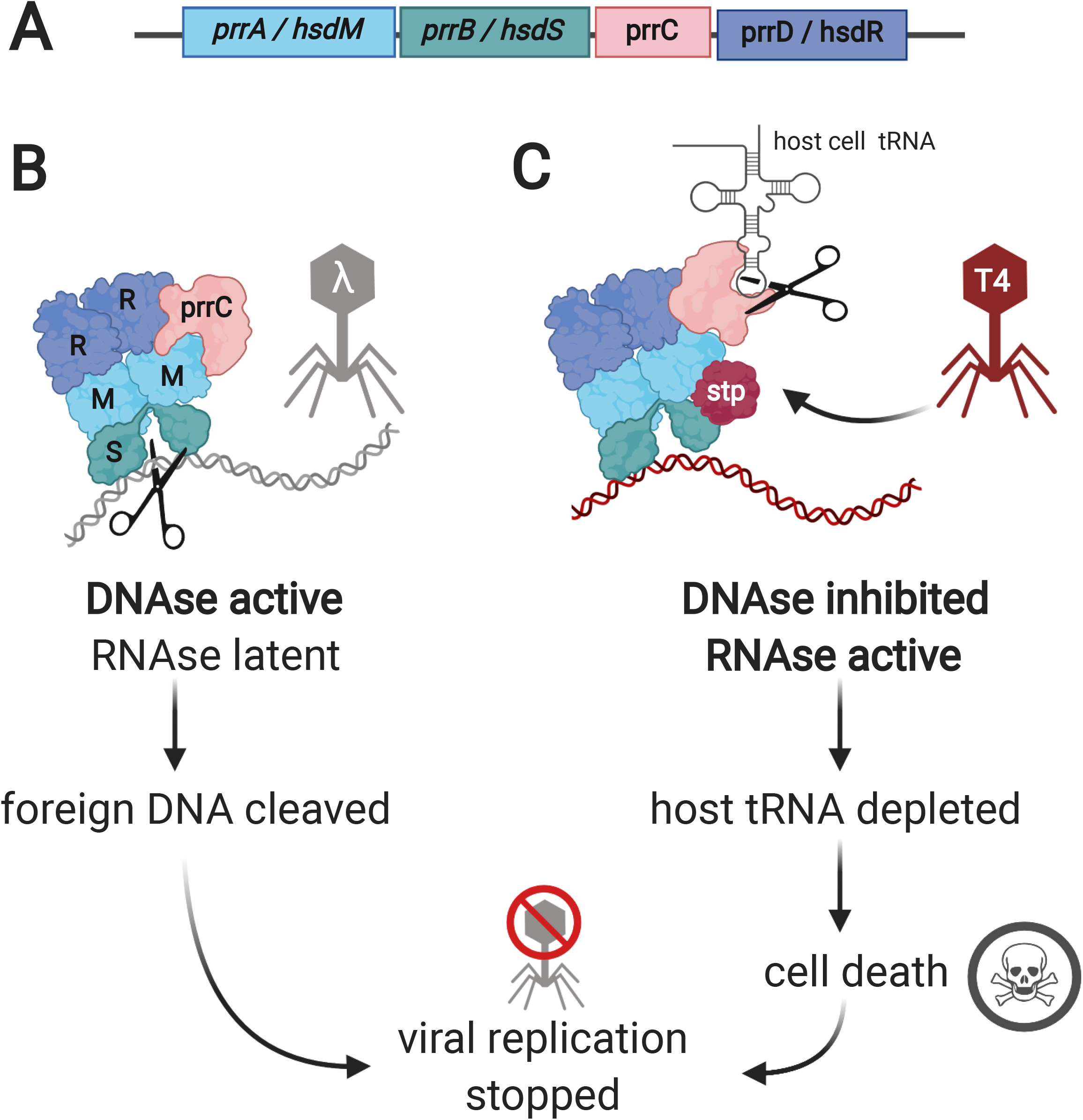
*E. colds EcoprrI* **(A)** provides an example of an RM system with two different avenues to halt the spread of a phage infection, either a classic DNAse-based Type I RM defense **(B)** or an RNAse-based abortive infection strategy **(C)**. Encoded by a four gene operon **(A)**, the complex consists of PrrC, an tRNA^Lys^-specific anticodon nuclease, associated with a typical Type I RM system (a methylase (PrrA / HsdM, “M”), a specificity determinant that interacts with the DNA binding site (PrrB / SsdS, “S”), and a restriction enzyme (PrrD / HsdR, “R”). **(B)** EcoprrI assembles as a single protein complex, like other Type I RM systems, but with the addition of a latent, inactive PrrC subunit. During normal growth or infection by a susceptible virus, such as phage lambda, EcoprrI operates as a typical Type I restriction enzyme, and halts viral replication by cleaving the DNA of the infecting phage. **(C)** T4-phage encodes a resistance mechanism: a short peptide (Stp) which can bind to EcoprrI and inhibit its endonuclease activity, likely due to a conformational change. However, this same Stp-induced conformational change activates the complex’s PrrC anticodon nuclease which cleaves host tRNA^Lys^. This RNAse activity depletes the host’s tRNA^Lys^ which inhibits protein synthesis and kills the host. This “abortive infection” strategy, where the host cell detects resistant phage and sacrifices itself, stops viral replication and minimizes the spread of phage to the host’s vulnerable clonal kin. We propose that the co-occurrence of Type I RM and other RNAse toxin genes in the *Thiohalocapsa* sp. PB-PSB1 genome could represent an analogous system, combining both RM and abortive infection phage defenses.

Resolving these complex features in bacterial genomes opens exciting frontiers for investigations of microbial consortia and gives us a lens that allows us to examine how ecological interactions – from symbioses to predation—shape bacterial evolution.

## Methods

### Sampling and library preparation

Pink berry aggregates were sampled in July 2011 from Little Sippewissett Salt Marsh, as described previously, and DNA was extracted using a modified phenol chloroform protocol (see Supplemental Methods). We created three distinct samples from which DNA was extracted: a very large aggregate ~9 mm in diameter (berry9), a pool of 13 aggregates 2-3 mm in diameter (s01), and a pool of 10 aggregates of similar size (s02). Transposase-based Illumina Nextera XT libraries were constructed for samples berry9 and s02. Sample berry9 was sequenced via Illumina MiSeq (1Gb of 250 bp paired end reads), while sample s02 was sequenced with both Illumina HiSeq (150PE) and a MiSeq run (250PE).

SMRTbell libraries for Pacific Biosciences sequence were constructed from 900 ng of berry9 DNA and ~1 microgram of s01 DNA. Sample s01 was size selected by Blue Pippin, while berry9 was selected with Ampure beads. In total, 42 SMRT cells were sequenced using PacBio RSII from these two libraries using a combination of P4C2 and P5C3 chemistries (25 cells from the berry9 Ampure library and 17 from the s01 BluePippin library). While Blue Pippin size-selection increased the proportion of reads greater than 8 kb, library sequence yield was poor when compared to the more robust Ampure bead library (44 Mb of filtered subreads per cell vs. 8.1 Gb of filtered subreads per cell). The PacBio data were pooled for further processing and are overwhelmingly represented by the sequence data from the berry9 sample (92% of filtered subread basepairs).

### Metagenomic assembly

Illumina sequence from sample s02 was trimmed and filtered with sga (preprocess -q 20 - f 20 -m 59 --pe-mode=1; Simpson and Durbin, 2012), adapter filtered with TagDust (Lassmann et al 2009), and assembled with idba_ud (maxk=250; v 1.0.9) (Peng et al 2012). This Illumina assembly was binned and curated as described previously (Wilbanks et al 2014). Binned sequence was reassembled and the MAGs were quality assessed with CheckM (Parks et al 2015).

PacBio sequence data were error corrected using SMRT Analysis 2.2, yielding 474 Mb of error corrected reads. Error corrected reads longer than 7 kb were assembled with the HGAP assembler (v. 3.3) using a reduced genome size parameter (genomeSize = 5,000,000) to increase tolerance of uneven coverage and an increased overlap error rate parameter (ovlErrorRate = 0.10) and overlap length (ovlMinLen =60) to encourage contig merging. The topology of the assembly graph (Celera Assembler’s “best.edges”) for the PacBio assembly was visualized in Gephi (Bastian et al 2009) to determine connectivity between fragmented contigs. This connectivity was used as an additional metric for binning validation, analogous to an approach proposed for short read assemblies (Mallawaarachchi et al 2020). Metagenomic contigs were quality checked and taxonomically identified as described in the Supplemental Methods.

### Methylation analysis and metagenomic binning

Methylated bases and their associated motifs were detected on the assembled contigs using the SMRT Analysis v. 2.2 module RS_Modification_and_Motif_Analysis.1 with an *in silico* control model (modification quality value > 20). For detected motifs, we computed the percentage of methylated motifs out of the total instances of that motif on each contig. The vector of percent methylations for all characteristic motifs represents the contig’s methylation profile. Contigs were hierarchically clustered in Cytoscape (v 3.5.1) using Clustermaker2 (Morris et al 2011) based on the Euclidean distance between square root transformed methylation profiles. t-distributed stochastic nearest neighbor clustering (t-SNE) of contigs based on both methylation profiles and tetranucleotide frequencies was performed with the Rtsne package (van der Maaten 2014) and visualized with ggplot2 (Wickham 2016) in R. Binned sequences were individually reassembled with HGAP. The PB-PSB1 MAG was circularized and manually curated using Geneious (v R11). MAGs were polished with pilon (Walker et al. 2014) using both Illumina and PacBio data, and corrections were manually verified for short-read mapping errors.

The methylated motifs in each MAG were predicted independently using SMRT Analysis (v. 2.2). For incomplete genomes (e.g. alphaproteobacterial MAGs), both PacBio- and Illumina-assembled versions of the MAG were used as the reference genome used to recruit the PacBio reads for methylation analysis. Restriction modification system annotation and motif matching was accomplished by comparison of the genome sequences and methylated sequence motifs with the Restriction Enzyme Database (REBASE) (Roberts et al 2015), as previously described (Blow et al 2016).

### Data availability

All sequence data has been deposited in DDBJ/ENA/GenBank under BioProject PRJNA684324. The accession numbers for the Short Read Archive and genome / metagenome data are provided in Supplemental Table 2.

## References

Albertsen M, Hugenholtz P, Skarshewski A, Nielsen KL, Tyson GW, Nielsen PH (2013). Genome sequences of rare, uncultured bacteria obtained by differential coverage binning of multiple metagenomes. Nature Biotechnology 31:533.

Banfield JF, Anantharaman K, Williams KH, Thomas BC (2017). Complete 4.55 megabase pair genome of “Candidates Fluviicola riflensis,” curated from short-read metagenomic sequences. Genome Announc.5:e01299–17

Bao YJ, Shapiro BJ, Lee SW, Ploplis VA, Castellino FJ (2016). Phenotypic differentiation of Streptococcus pyogenes populations is induced by recombination-driven gene-specific sweeps. Scientific Reports 6: 36644.

Bartlett DH, Wright ME, Silverman M (1988). Variable expression of extracellular polysaccharide in the marine bacterium *Pseudomonas atlantica* is controlled by genome rearrangement. PNAS 85:3923–3927.

Bastian M, Heymann S, Jacomy M: Gephi: an open source software for exploring and manipulating networks. International AAAI Conference on Weblogs and Social Media. 2009.

Beaulaurier J, Zhu SJ, Deikus G, Mogno I, Zhang XS, Davis-Richardson A et al (2018). Metagenomic binning and association of plasmids with bacterial host genomes using DNA methylation. Nature Biotechnology 36:61.

Beaulaurier J, Schadt EE, Fang G (2019). Deciphering bacterial epigenomes using modern sequencing technologies. Nat Rev Genet 20:157–172.

Beitel CW, Froenicke L, Lang JM, Korf IF, Michelmore RW, Eisen JA et al (2014). Strain-and plasmid-level deconvolution of a synthetic metagenome by sequencing proximity ligation products. Peerj 2.

Berube PM, Rasmussen A, Braakman R, Stepanauskas R, Chisholm SW (2019). Emergence of trait variability through the lens of nitrogen assimilation in Prochlorococcus. eLlife 8:e41043.

Blow MJ, Clark TA, Daum CG, Deutschbauer AM, Fomenkov A, Fries R et al (2016). The epigenomic landscape of prokaryotes. Plos Genetics 12:e1005854.

Buryanov Y, Shevchuk T (2005). The use of prokaryotic DNA methyltransferases as experimental and analytical tools in modern biology. Anal Biochem 338:1–11.

Chen LX, Anantharaman K, Shaiber A, Eren AM, Banfield JF (2020). Accurate and complete genomes from metagenomes. Genome Research 30:315–333.

Dick GJ, Andersson AF, Baker BJ, Simmons SL, Thomas BC, Yelton aP et al (2009). Community-wide analysis of microbial genome sequence signatures. Genome biology 10:R85–R85.

Doré H, Farrant GK, Guyet U, Haguait J, Humily F, Ratin M et al (2020). Evolutionary mechanisms of long-term genome diversification associated with niche partitioning in marine picocyanobacteria. Frontiers in Microbiology 11:e567431.

Everitt RG, Didelot X, Batty EM, Miller RR, Knox K, Young BC et al (2014). Mobile elements drive recombination hotspots in the core genome of *Staphylococcus aureus*. Nature Communications 5: 3956.

Hehemann JH, Arevalo P, Datta MS, Yu XQ, Corzett CH, Henschel A et al (2016). Adaptive radiation by waves of gene transfer leads to fine-scale resource partitioning in marine microbes. Nature Communications 7: 12860.

Hewson I, Poretsky RS, Dyhrman ST, Zielinski B, White AE, Tripp HJ et al (2009). Microbial community gene expression within colonies of the diazotroph, *Trichodesmium*, from the Southwest Pacific Ocean. ISME J 3:1286–1300.

Higgins BP, Carpenter CD, Karls AC (2007). Chromosomal context directs high-frequency precise excision of IS492 in Pseudoalteromonas atlantica. PNAS 104:1901–1906.

Huson DH, Auch AF, Qi J, Schuster SC (2007). MEGAN analysis of metagenomic data. Genome Research 17:377–386.

Kaneko T, Nakajima N, Okamoto S, Suzuki I, Tanabe Y, Tamaoki M et al (2007). Complete genomic structure of the bloom-forming toxic cyanobacterium *Microcystis aeruginosa* NIES-843. DNA Research 14:247–256.

Kaufmann G (2000). Anticodon nucleases. Trends Biochem Sci 25:70–74.

Koonin EV, Makarova KS, Wolf YI (2017). Evolutionary genomics of defense systems in *Archaea* and Bacteria. Annual Review of Microbiology 71:233–261.

Lassmann T, Hayashizaki Y, Daub CO (2009). TagDust-a program to eliminate artifacts from next generation sequencing data. Bioinformatics 25:2839–2840.

Maguire F, Jia BF, Gray KL, Lau WYV, Beiko RG, Brinkman FSL (2020). Metagenome-assembled genome binning methods with short reads disproportionately fail for plasmids and genomic islands. Microb Genomics 6. 10.1099/mgen.0.000436.

Mallawaarachchi V, Wickramarachchi A, Lin Y (2020). GraphBin: refined binning of metagenomic contigs using assembly graphs. Bioinformatics 36:3307–3313.

Meyer F, Hofmann P, Belmann P, Garrido-Oter R, Fritz A, Sczyrba A et al (2018). AMBER: Assessment of Metagenome BinnERs. Gigascience 7: giy069.

Morris JH, Apeltsin L, Newman AM, Baumbach J, Wittkop T, Su G et al (2011). clusterMaker: a multi-algorithm clustering plugin for Cytoscape. BMC Bioinformatics 12: 436

Murray NE (2000). Type I restriction systems: Sophisticated molecular machines (a legacy of Bertani and Weigle). Microbiology and Molecular Biology Reviews 64:412–434.

Newton ILG, Bordenstein SR (2011). Correlations Between Bacterial Ecology and Mobile DNA. Curr Microbiol 62:198–208.

Oliveira PH, Touchon M, Cury J, Rocha EPC (2017). The chromosomal organization of horizontal gene transfer in bacteria. Nature Communications 8: 841.

Olson ND, Treangen TJ, Hill CM, Cepeda-Espinoza V, Ghurye J, Koren S et al (2019). Metagenomic assembly through the lens of validation: recent advances in assessing and improving the quality of genomes assembled from metagenomes. Brief Bioinform 20:1140–1150.

Parks DH, Imelfort M, Skennerton CT, Hugenholtz P, Tyson GW (2015). CheckM: assessing the quality of microbial genomes recovered from isolates, single cells, and metagenomes. Genome Research 25:1043–1055.

Peng Y, Leung HCM, Yiu SM, Chin FYL (2012). IDBA-UD: a de novo assembler for single-cell and metagenomic sequencing data with highly uneven depth. Bioinformatics 28:1420–1428.

Roberts RJ, Vincze T, Posfai J, Macelis D (2015). REBASE-a database for DNA restriction and modification: enzymes, genes and genomes. Nucleic acids research 43:D298–D299.

Rocap G, Larimer FW, Lamerdin J, Malfatti S, Chain P, Ahlgren NA et al (2003). Genome divergence in two Prochlorococcus ecotypes reflects oceanic niche differentiation. Nature 424:1042–1047.

Sanchez-Romero MA, Casadesus J (2020). The bacterial epigenome. Nature Reviews Microbiology 18:7–20.

Sieber CMK, Probst AJ, Sharrar A, Thomas BC, Hess M, Tringe SG et al (2018). Recovery of genomes from metagenomes via a dereplication, aggregation and scoring strategy. Nature Microbiology 3:836.

Stewart RD, Auffret MD, Warr A, Wiser AH, Press MO, Langford KW et al (2018). Assembly of 913 microbial genomes from metagenomic sequencing of the cow rumen. Nature Communications 9:870.

Touchon M, Rocha EPC (2007). Causes of insertion sequences abundance in prokaryotic genomes. Molecular Biology and Evolution 24:969–981.

Tyndall C, Meister J, Bickle TA (1994). The *Escherichia coli* Prr region encodes a functional type-Ic DNA restriction system closely integrated with an anticodon nuclease gene. Journal of Molecular Biology 237:266–274.

Tyson GW, Chapman J, Hugenholtz P, Allen EE, Ram RJ, Richardson PM et al (2004). Community structure and metabolism through reconstruction of microbial genomes from the environment. Nature 428:37–43.

van der Maaten L (2014). Accelerating t-SNE using Tree-Based Algorithms. J Mach Learn Res 15:3221–3245.

White RA, Bottos EM, Chowdhury TR, Zucker JD, Brislawn CJ, Nicora CD et al (2016). Moleculo long-read sequencing facilitates assembly and genomic binning from complex soil metagenomes. Msystems 1:e00045–00016.

Wickham H (2016). ggplot2: Elegant Graphics for Data Analysis. Springer-Verlag: New York.

Wilbanks EG, Jaekel U, Salman V, Humphrey PT, Eisen JA, Faccioti MT et al (2014). Microscale sulfur cycling in the phototrophic pink berry consortia of the Sippewissett Salt Marsh. Environmental Microbiology 16:3398–3415.

Ziebuhr W, Krimmer V, Rachid S, Lossner I, Gotz F, Hacker J (1999). A novel mechanism of phase variation of virulence in Staphylococcus epidermidis: evidence for control of the polysaccharide intercellular adhesin synthesis by alternating insertion and excision of the insertion sequence element IS256. Molecular Microbiology 32:345–356.

