## Supplemental Figures Tables and Methods for "Metagenomic methylation patterns resolve complex microbial genomes"

**Supplemental Methods**
*Sampling and DNA extraction*

"Pink berries" were collected from the sediment-water interface of a shallow marsh pool of the Little Sippewissett Salt Marsh, Falmouth, MA USA (41°34'33.01"N, 70°38'21.24"W) and washed with 0.2 micron filtered marsh water. We created 3 distinct samples from which DNA was extracted: a very large aggregate ~9 mm in diameter (berry9), a pool of 13 aggregates 2-3 mm in diameter (s01), a pool of 10 aggregates of similar size (s02). Berries in each sample were chemically disaggregated by 1 hour incubation at 37 ºC in a 5 M urea solution, pelleted, and washed 3x in 1x PBS in a 1.5 mL tube. High molecular weight DNA was extracted using a phenol chloroform protocol based on that of Dojka et al (1998) omitting all bead beating and vigorous mixing. Briefly, samples were resuspended in 500 uL of 2x buffer A (200 mM Tris pH 8, 50 mM EDTA, 200 mM NaCl, 2 mM sodium citrate dihydrate, 10 mM $CaCl_2$ dihydrate) with lysozyme (3 mg/mL), gently homogenized with a pestle and then incubated for 40 minutes at 37 ºC. Samples were gently inverted, and then proteinase K (to 1.2 mg/mL) and sodium dodecyl sulfate (10 uL of 20% w/v, SDS) were added, the tubes gently inverted, and the mixture was incubated for 30 min at 50°C. 500 uL of phenol-chloroform-isoamyl alcohol (phenol : $CHCl_3$ : IAA; 24:24:1) and 60 uL of 20% (w/v) SDS were added, placed on a rotating platform for 5 minutes to emulsify, and then samples were spun at 12xkg for 20 minutes at 4 ºC. Supernatant was re-extracted with 1 volume phenol:$CHCl_3$:IAA, and then DNA was precipitated with 1 volume isopropanol and 0.1 volume sodium acetate (3M, pH 5.2) by incubation on ice for 20 minutes followed by 20 minutes spinning at 12xkg at 4 ºC. Pellets were rinsed in 70% cold ethanol, and resuspended gently in nuclease free water.

*Taxonomic identification, annotation and QC*
Open reading frames on every contig were predicted with prokka (Seemann 2014) searched against to NR using BLASTP (v 2.2.25) (Altschul et al 1997) and taxonomically identified using MEGAN (Huson et al 2011). Taxonomic classification of contigs was based on conserved marker genes where present (Wu et al 2013), or else from the consensus of taxonomic assignments from all open reading frames. Where there was no clear taxonomic consensus or if majority of matches were to viral sequences, contigs were labeled "Unclassified" (grey fill, Figure 2). CheckM was used to assess marker gene based completeness and contamination of methylation-based bins (Parks et al 2015). Annotation of MAGs was performed with PGAP (Tatusova et al 2016), ISEScan (Xie and Tang 2017), PhiSpy (Akhter et al 2012), and CARD RGI (Alcock et al 2020). Whole genome alignments were conducted with progressiveMauve (Darling et al 2010).

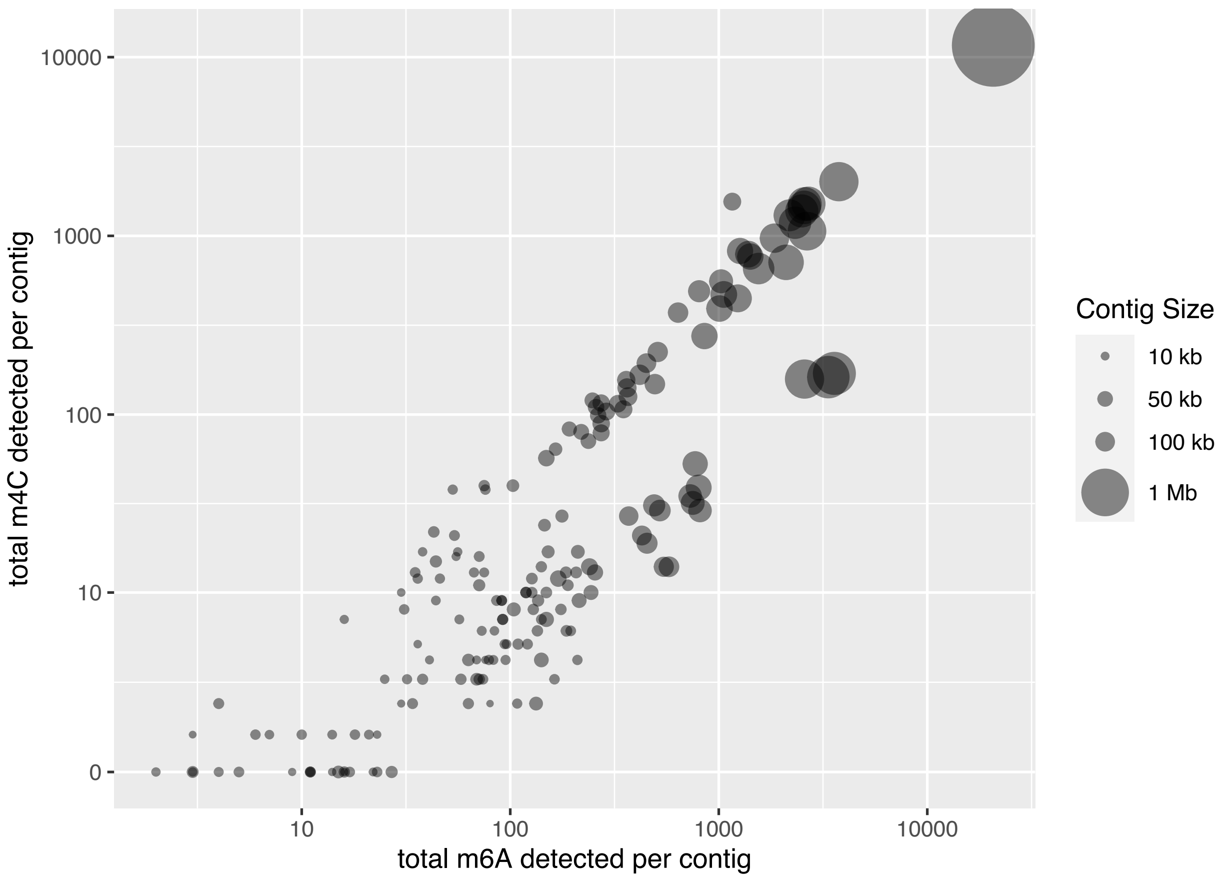


**Supplementary Figure 1**. The total number of $N^6$ -methyladenine (6mA) and $N^4$ -methylcytosine (4mC) detected on each contig in the assembly. Point size is scaled to represent the length of the contig sequence. Methylation detection thresholds were defined as a modification QV $\geq 20$, *i.e.* p-value $\leq 0.01$).

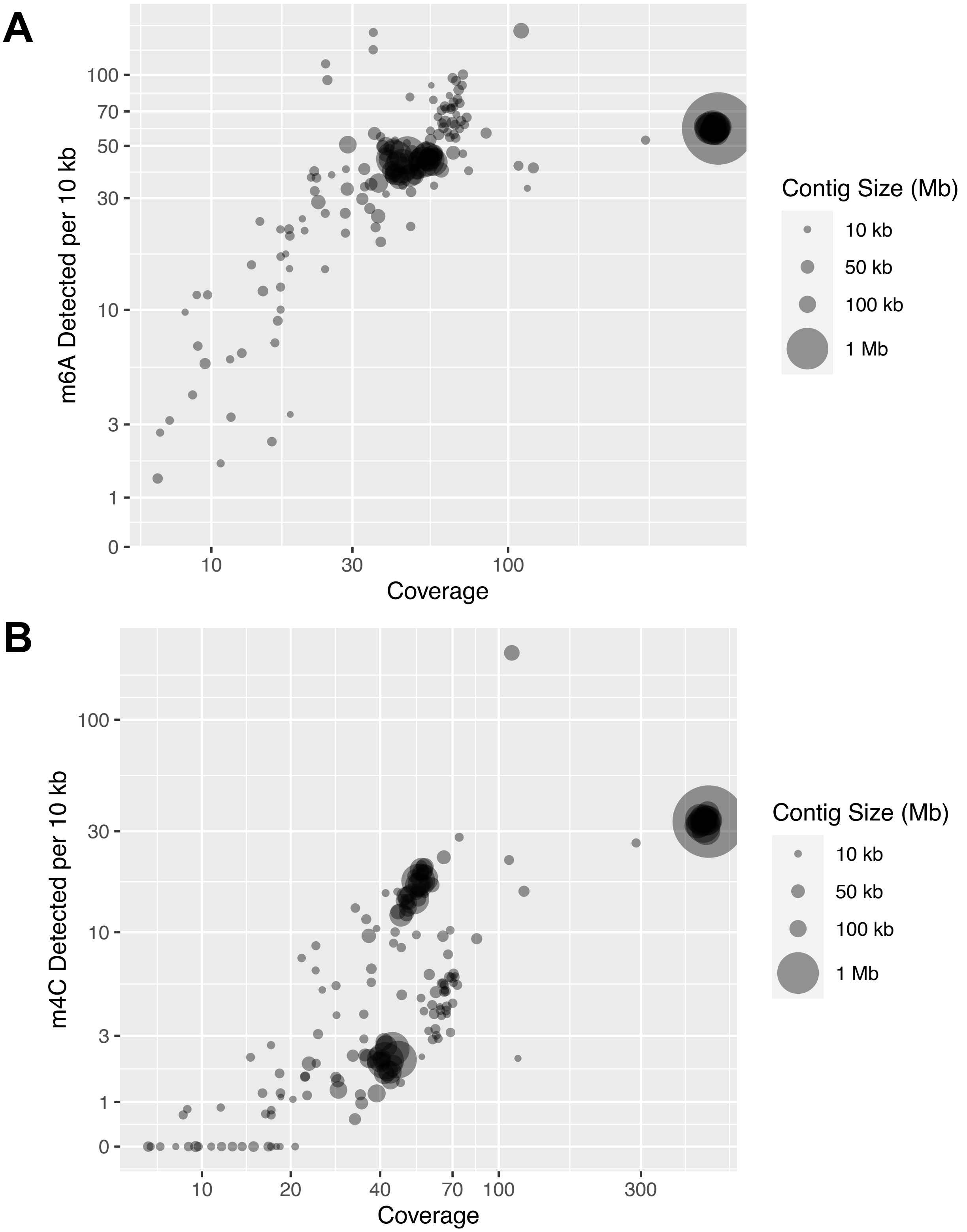


**Supplementary Figure 2.** The number A) m6A and (B) m4C modifications per 10 kb of assembled sequence on each contig. Point size is scaled to represent the length of the contig sequence. Methylation detection thresholds were defined as a modification QV $\geq$ 20, *i.e.* p-value $\leq$ 0.01).

**Supplemental Table 1.** 6-mA and 4-mC modified sequence motifs identified from large contigs in the assembled pacbio data using the SMRT Analysis workflow. The modification profiles on each contig for these 32 motifs, m01 – m32, are shown in Figure 1 (names correspond to those shown in Figure 1).

| motif ID | motif sequence |
|---|---|
| m01 | GAGNNNNNNAATC |
| m02 | GATTNNNNNNCTC |
| m03 | CTGCAG |
| m04 | CTCGAG |
| m05 | CTACKAC |
| m06 | AAGCTT |
| m07 | CCTNAGG |
| m08 | CCCACA |
| m09 | ACCCAG |
| m10 | GCGCGAT |
| m11 | GAGATG |
| m12 | AGGGCC |
| m13 | CCANNNNNNGTCA |
| m14 | TGACNNNNNNTGG |
| m15 | AGGCCT |
| m16 | GCANNNNNNTCAC |
| m17 | GTGANNNNNNTGC |
| m18 | AYGCCGC |
| m19 | GGATCC |
| m20 | GCTGAT |
| m21 | GCANNNNNNNGTTG |
| m22 | CAACNNNNNNNTGC |
| m23 | CGCGA |
| m24 | CGCANNNNNNGGG |
| m25 | CCAGCG |
| m26 | GGWCC |
| m27 | GAYCCC |
| m28 | RGATCY |
| m29 | GANTC |
| m30 | CANCATC |
| m31 | GCCAGG |
| m32 | GATGGA |

**Supplementary Figure 3.** t-SNE clustering of contigs based on methylation profiles. Equivalent to maintext figure 2A but colored by % GC content of the contig. In red are the 3 low coverage, low GC (<45%) contigs within methylation cluster 4 that were assigned to *Bacteroidetes* (unitig_260, unitig_246) and a putative *Bacteroidetes* prophage (unitig_174).

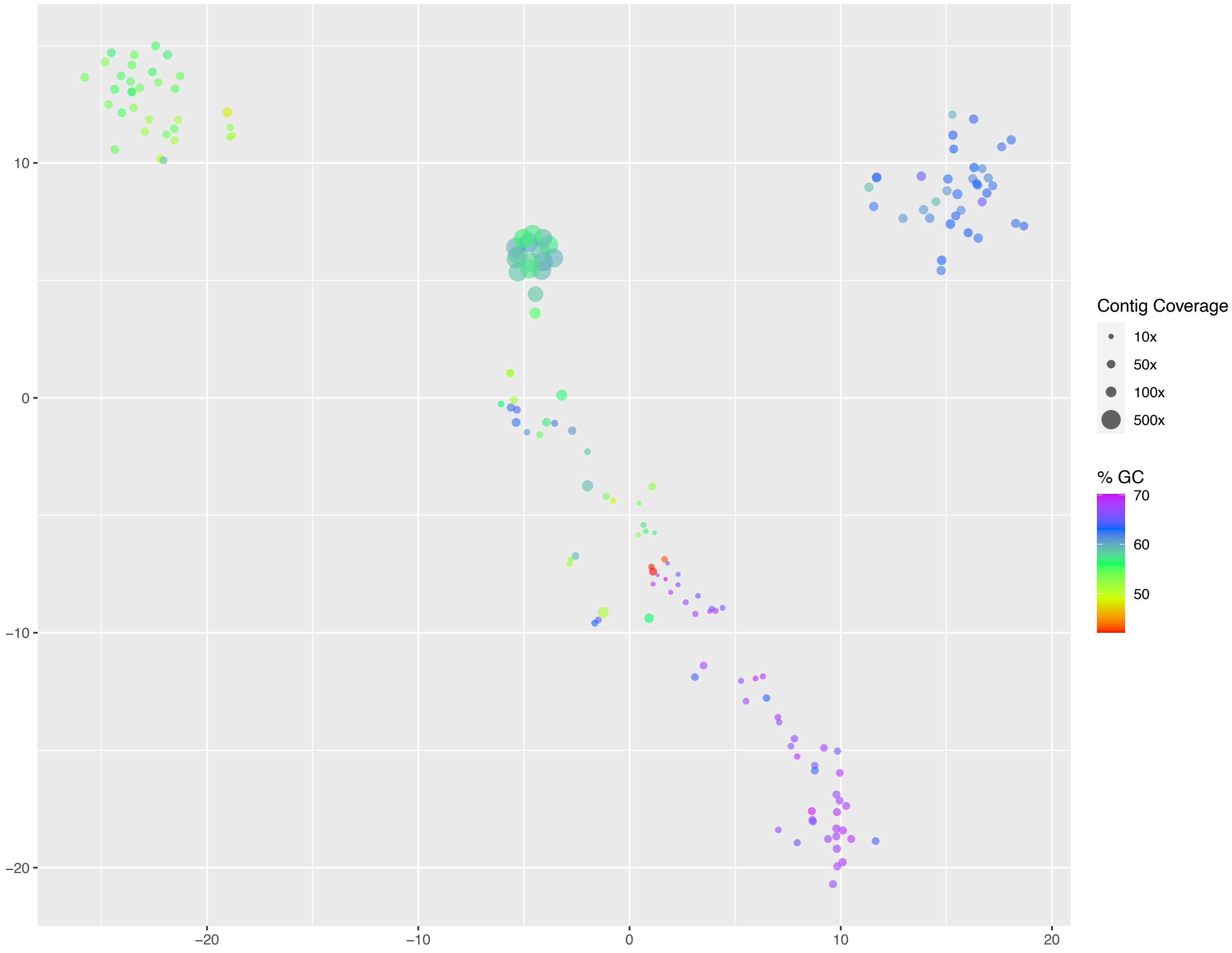

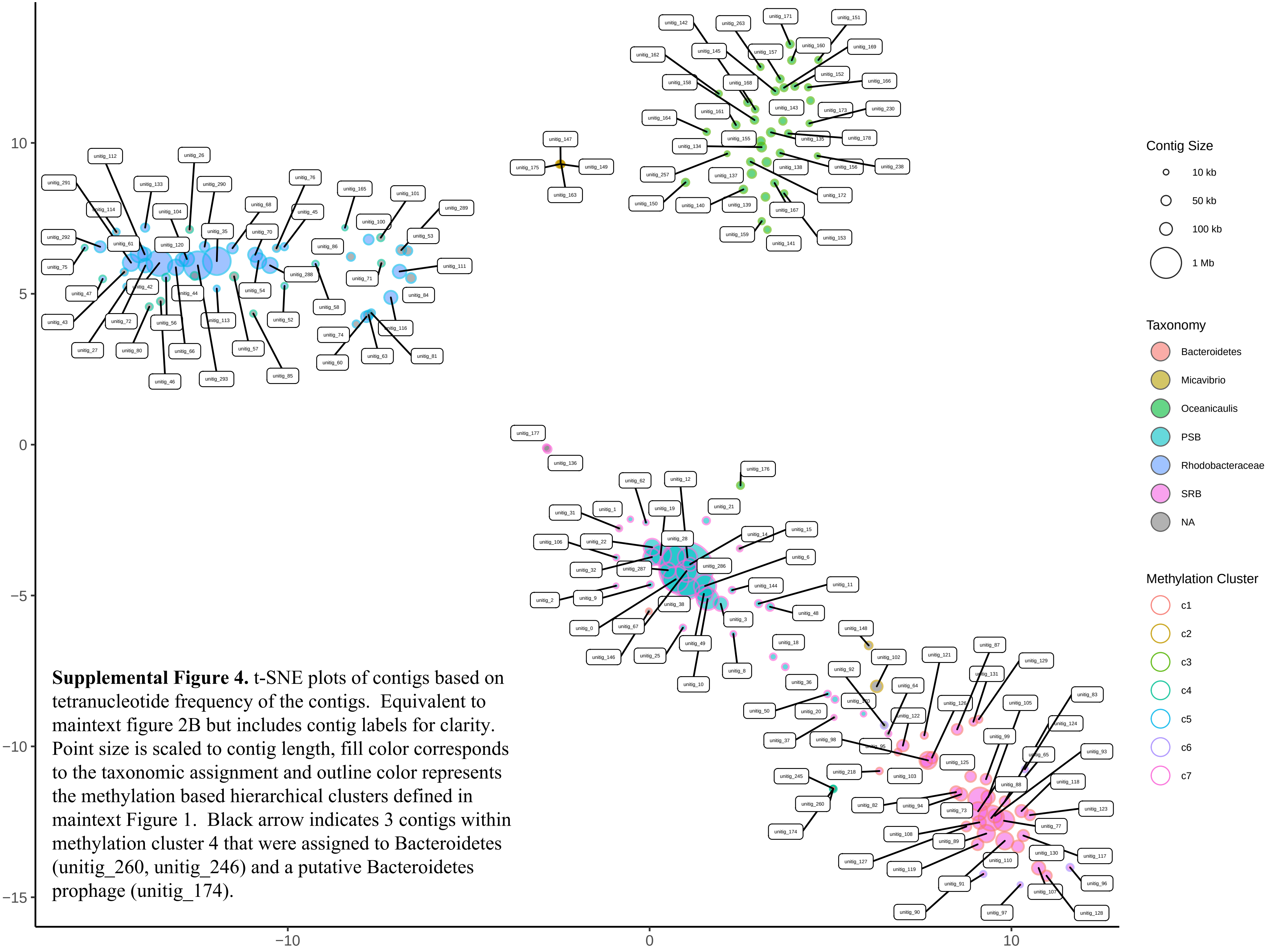


**Supplemental Figure 4.** t-SNE plots of contigs based on tetranucleotide frequency of the contigs. Equivalent to maintext figure 2B but includes contig labels for clarity. Point size is scaled to contig length, fill color corresponds to the taxonomic assignment and outline color represents the methylation based hierarchical clusters defined in maintext Figure 1. Black arrow indicates 3 contigs within methylation cluster 4 that were assigned to Bacteroidetes (unitig_260, unitig_246) and a putative Bacteroidetes prophage (unitig_174).
