## Supplemental Data 1 for "Metagenomic methylation patterns resolve complex microbial genomes"

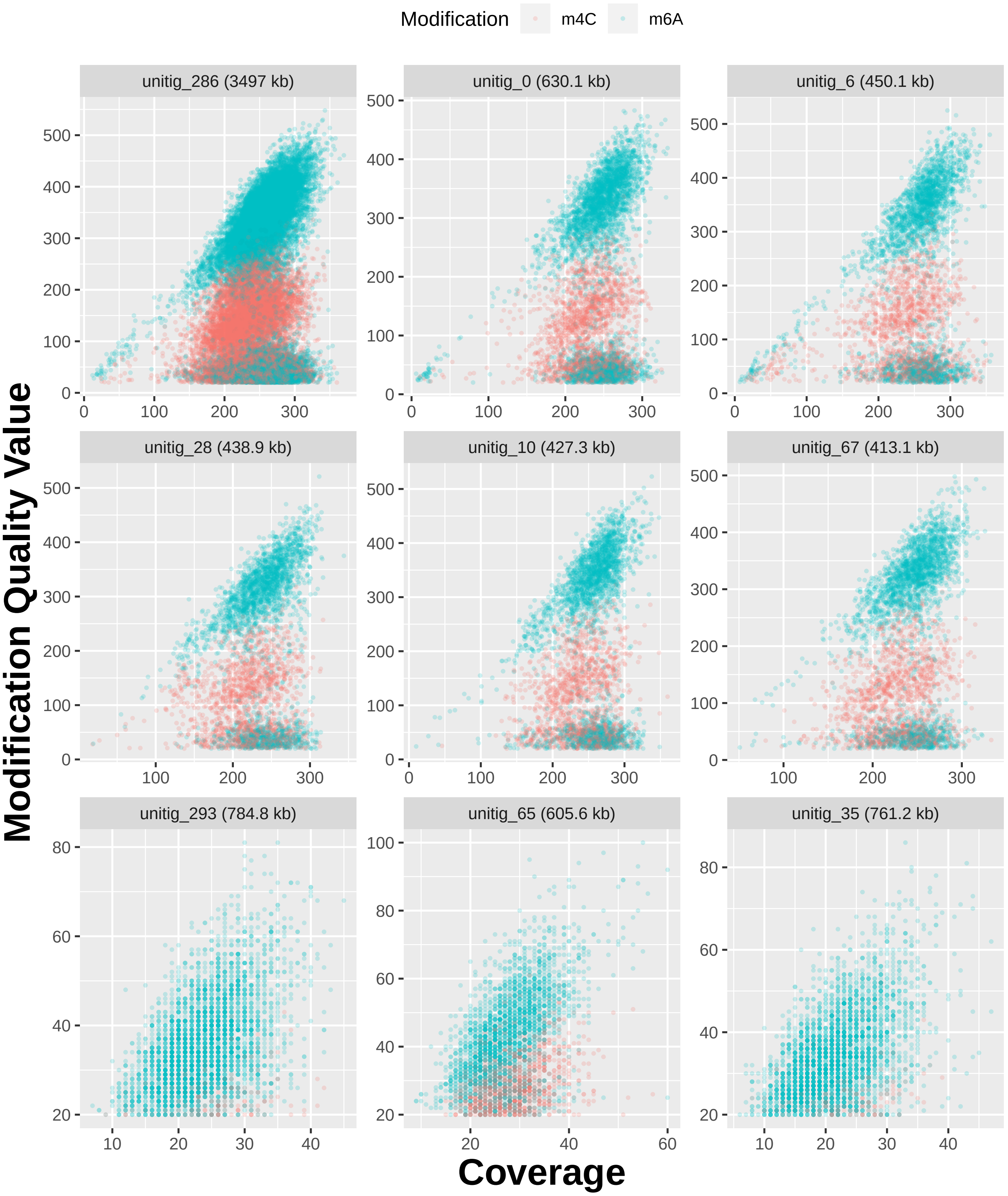

Modification
m4C
m6A
unitig_286 (3497 kb)
unitig_0 (630.1 kb)
unitig_6 (450.1 kb)
unitig_28 (438.9 kb)
unitig_10 (427.3 kb)
unitig_67 (413.1 kb)
unitig_293 (784.8 kb)
unitig_65 (605.6 kb)
unitig_35 (761.2 kb)
Modification Quality Value
Coverage

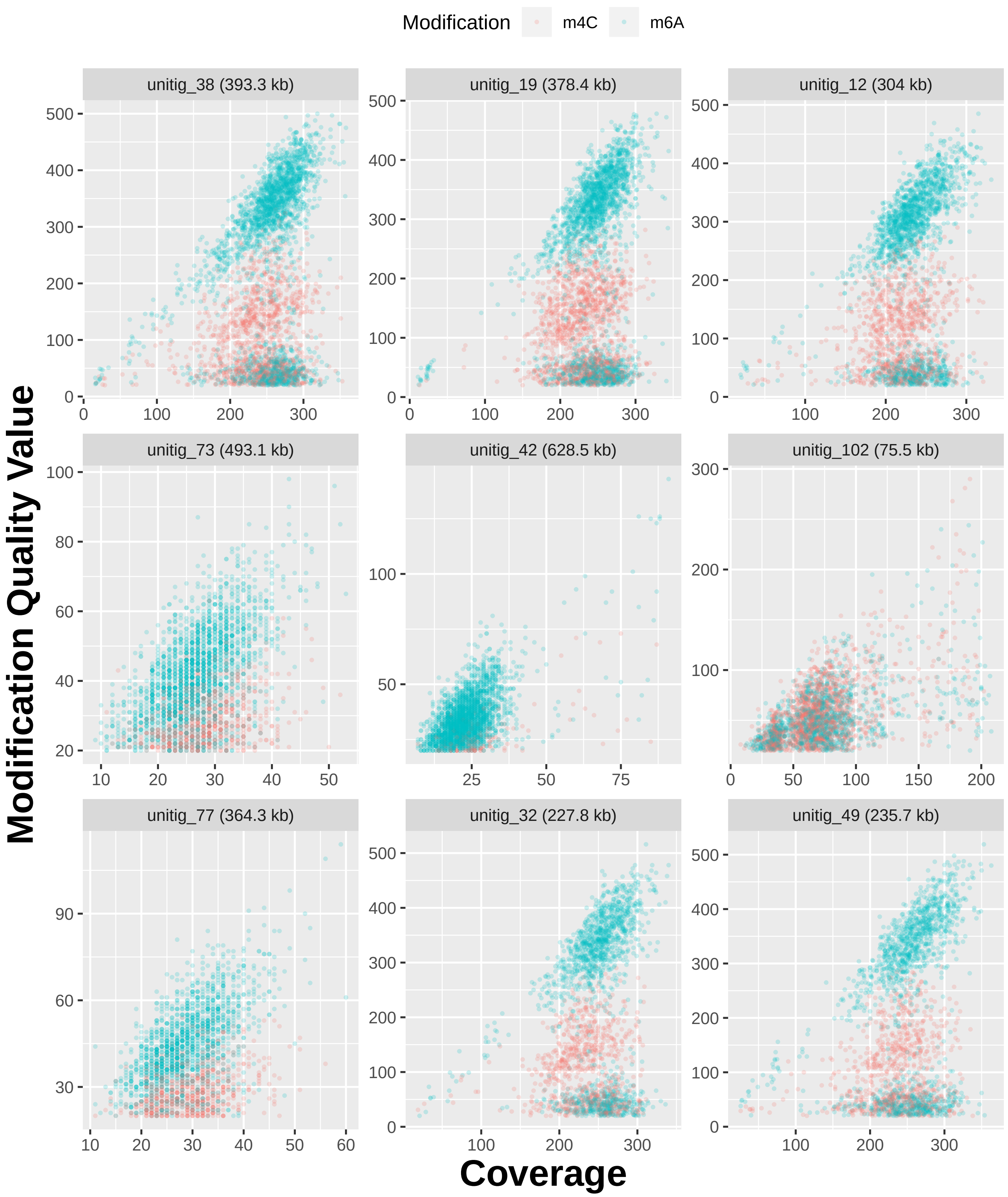

Modification
m4C
m6A
unitig_38 (393.3 kb)
unitig_19 (378.4 kb)
unitig_12 (304 kb)
unitig_73 (493.1 kb)
unitig_42 (628.5 kb)
unitig_102 (75.5 kb)
unitig_77 (364.3 kb)
unitig_32 (227.8 kb)
unitig_49 (235.7 kb)
Modification Quality Value
Coverage

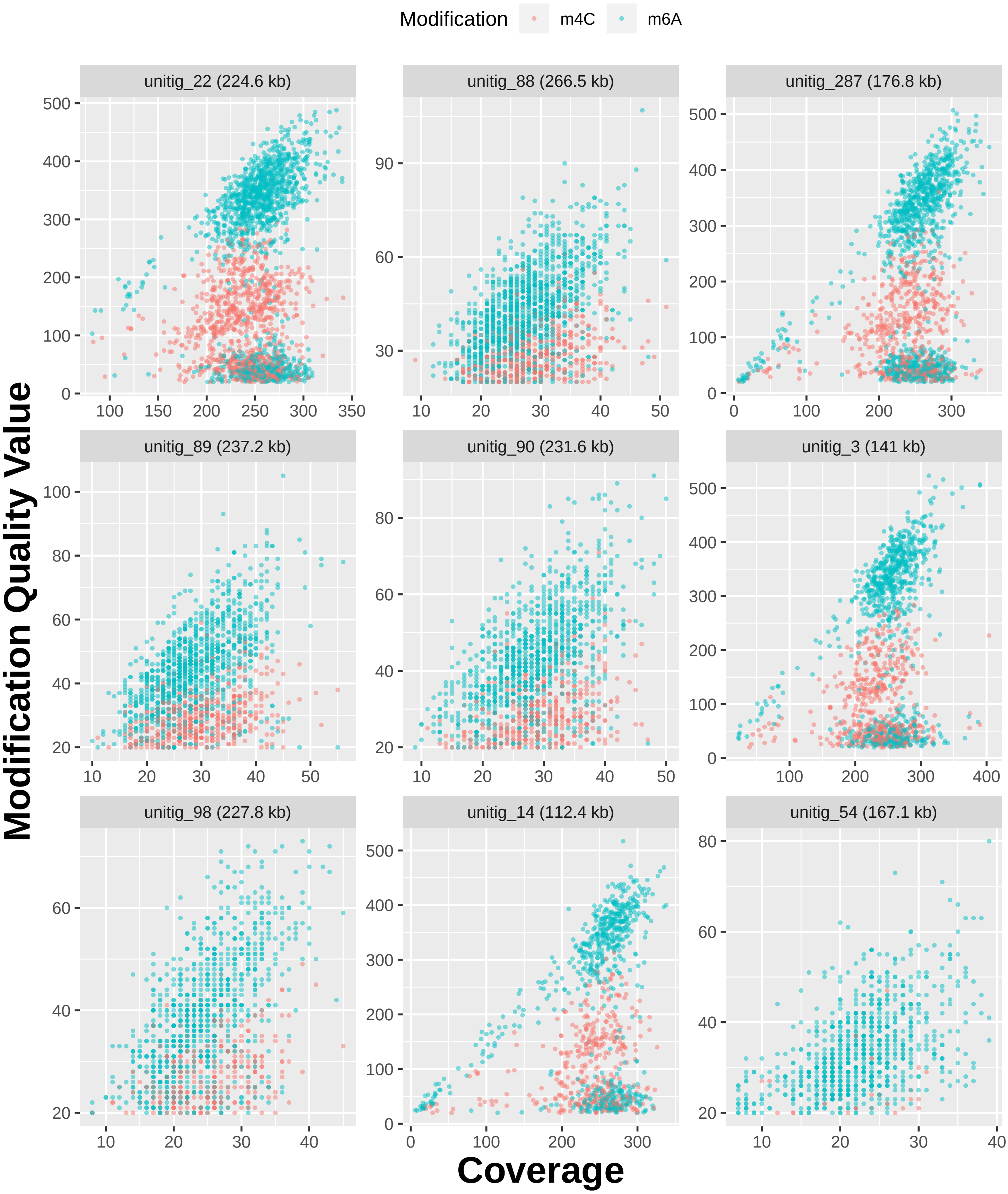
Modification
m4C
m6A
unitig_22 (224.6 kb)
unitig_88 (266.5 kb)
unitig_287 (176.8 kb)
unitig_89 (237.2 kb)
unitig_90 (231.6 kb)
unitig_3 (141 kb)
unitig_98 (227.8 kb)
unitig_14 (112.4 kb)
unitig_54 (167.1 kb)
Modification Quality Value
Coverage

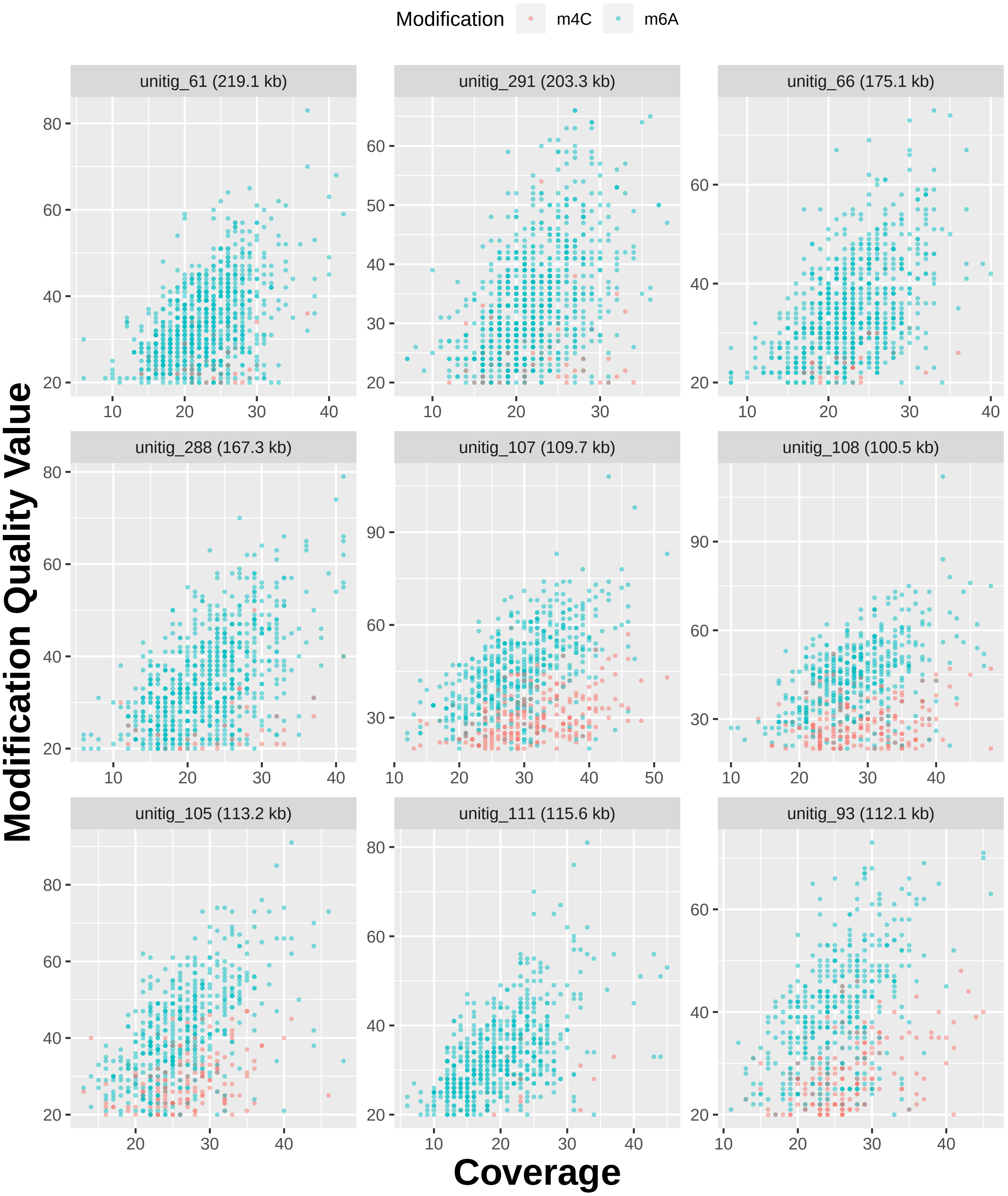

Modification
m4C
m6A
unitig_61 (219.1 kb)
unitig_291 (203.3 kb)
unitig_66 (175.1 kb)
unitig_288 (167.3 kb)
unitig_107 (109.7 kb)
unitig_108 (100.5 kb)
unitig_105 (113.2 kb)
unitig_111 (115.6 kb)
unitig_93 (112.1 kb)
Modification Quality Value
Coverage

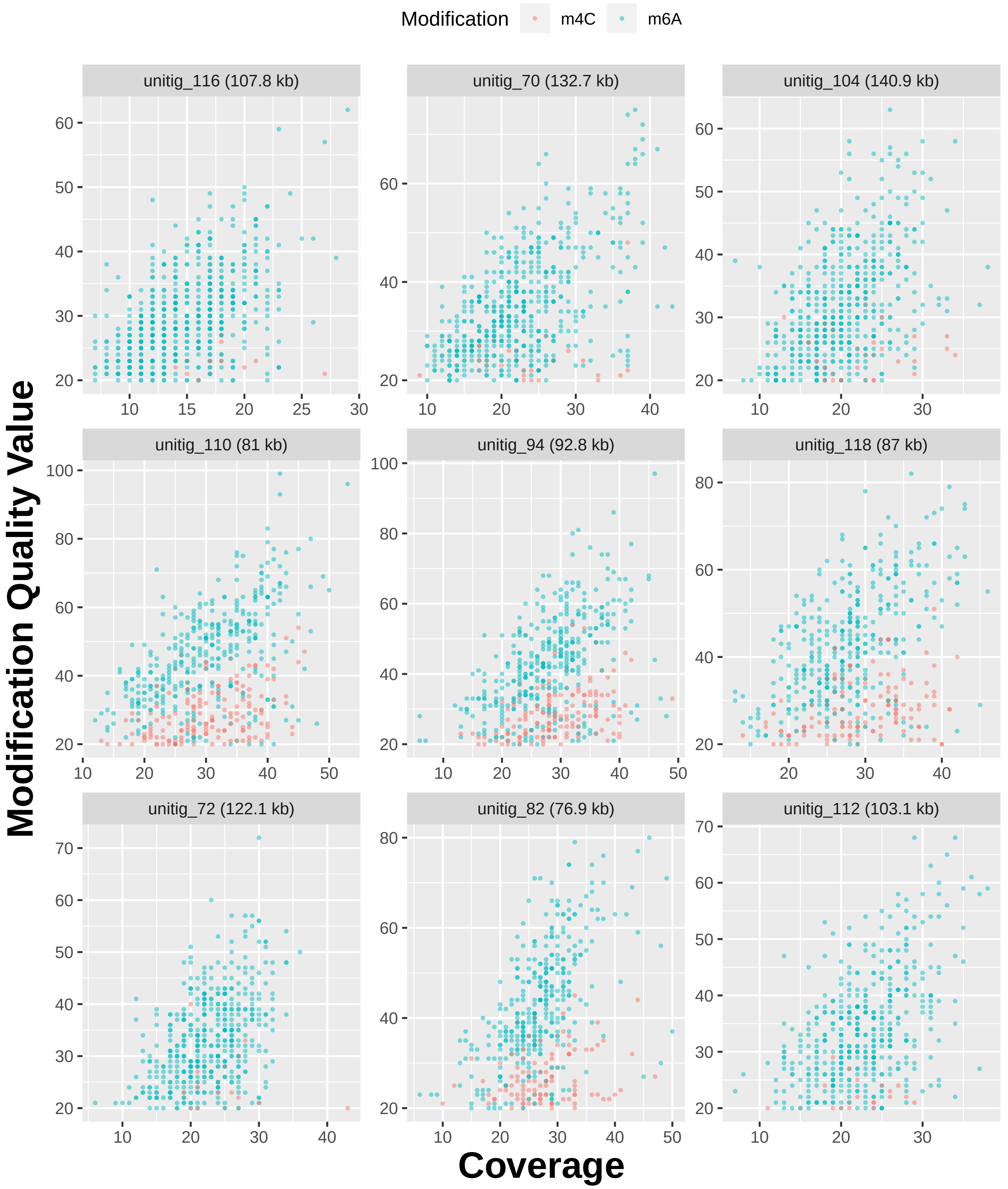

Modification
m4C
m6A
unitig_116 (107.8 kb)
unitig_70 (132.7 kb)
unitig_104 (140.9 kb)
unitig_110 (81 kb)
unitig_94 (92.8 kb)
unitig_118 (87 kb)
unitig_72 (122.1 kb)
unitig_82 (76.9 kb)
unitig_112 (103.1 kb)
Modification Quality Value
Coverage

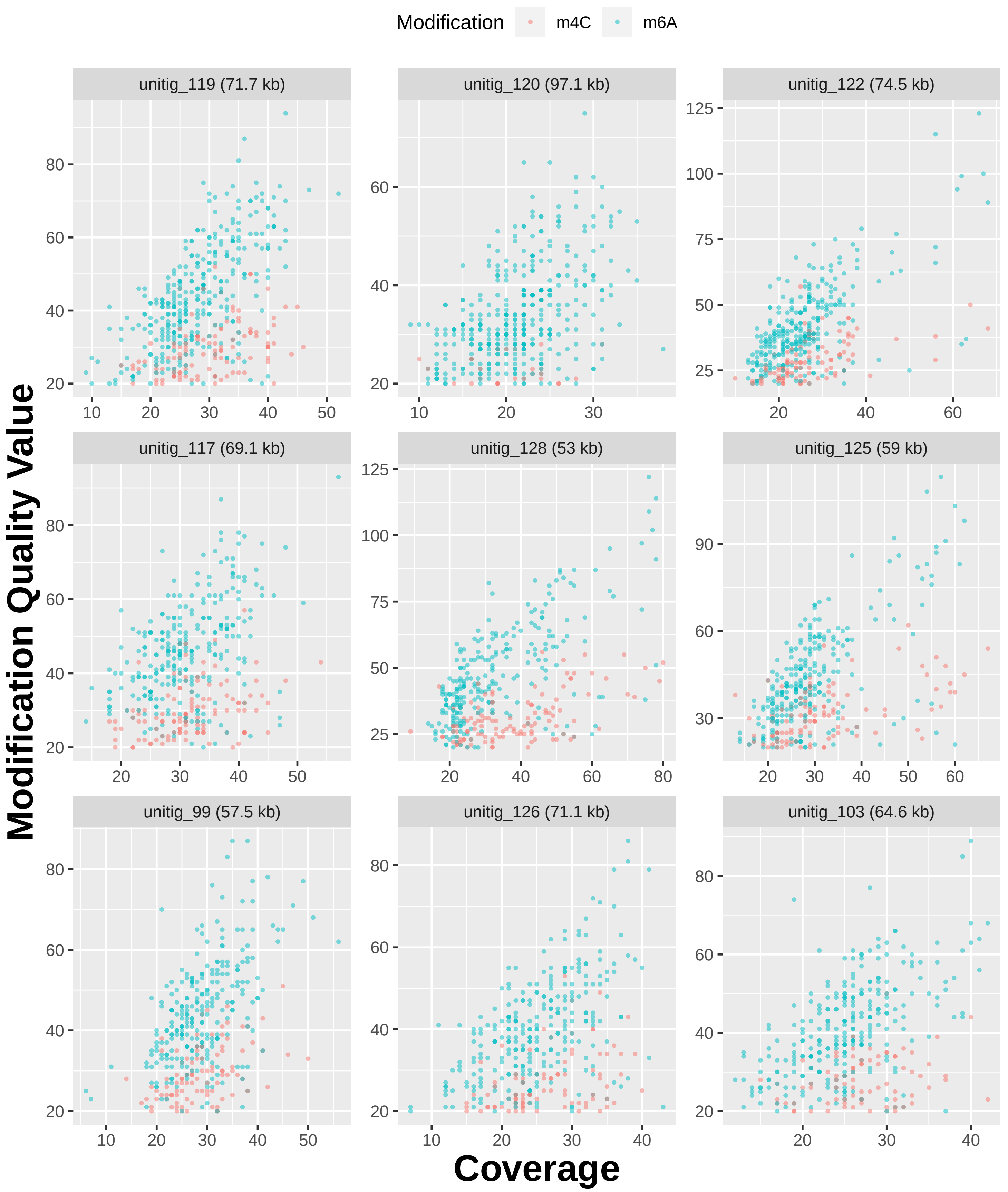

Modification
m4C
m6A
unitig_119 (71.7 kb)
unitig_120 (97.1 kb)
unitig_122 (74.5 kb)
unitig_117 (69.1 kb)
unitig_128 (53 kb)
unitig_125 (59 kb)
unitig_99 (57.5 kb)
unitig_126 (71.1 kb)
unitig_103 (64.6 kb)
Modification Quality Value
Coverage

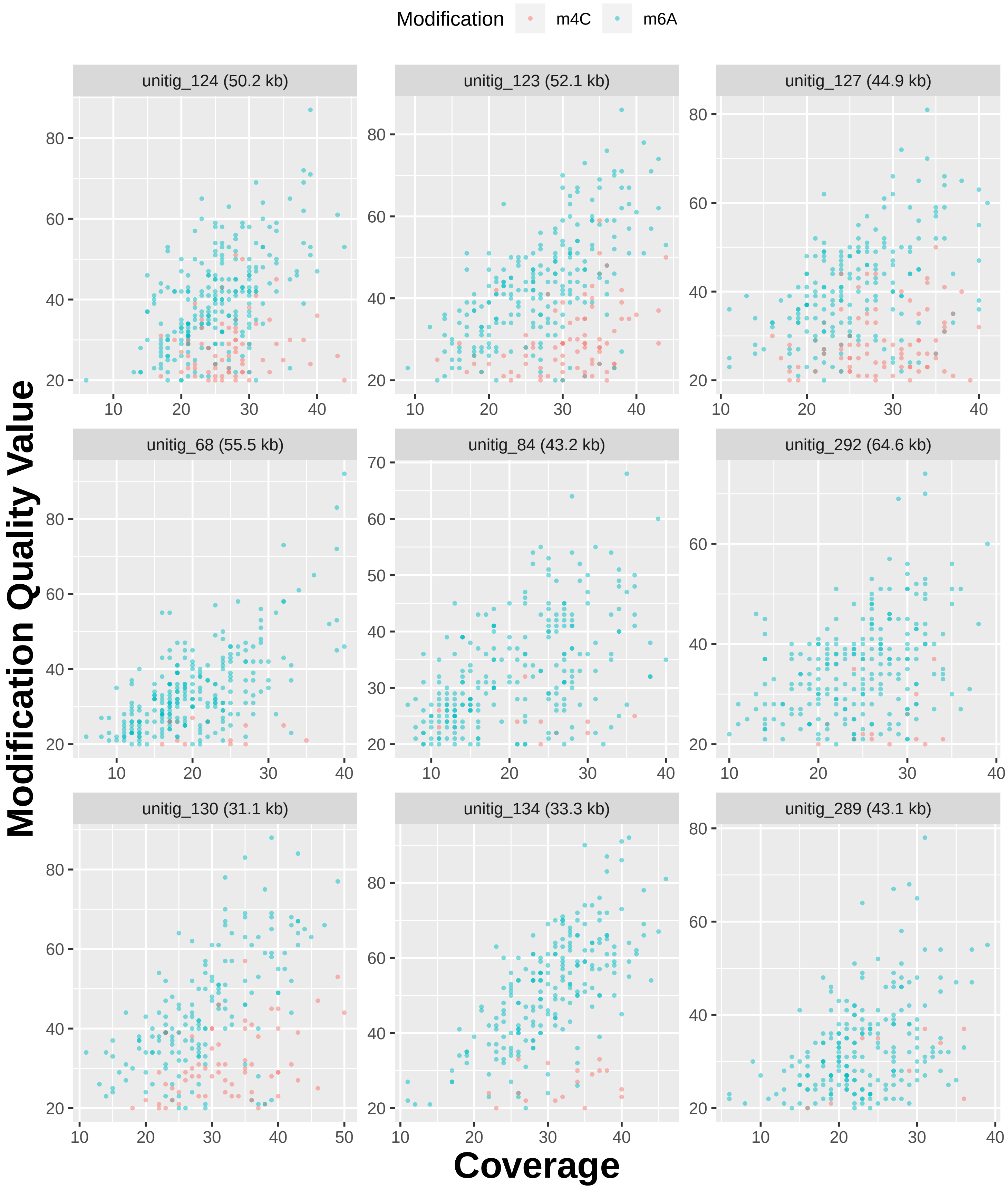

Modification
m4C
m6A
unitig_124 (50.2 kb)
unitig_123 (52.1 kb)
unitig_127 (44.9 kb)
unitig_68 (55.5 kb)
unitig_84 (43.2 kb)
unitig_292 (64.6 kb)
unitig_130 (31.1 kb)
unitig_134 (33.3 kb)
unitig_289 (43.1 kb)
Modification Quality Value
Coverage

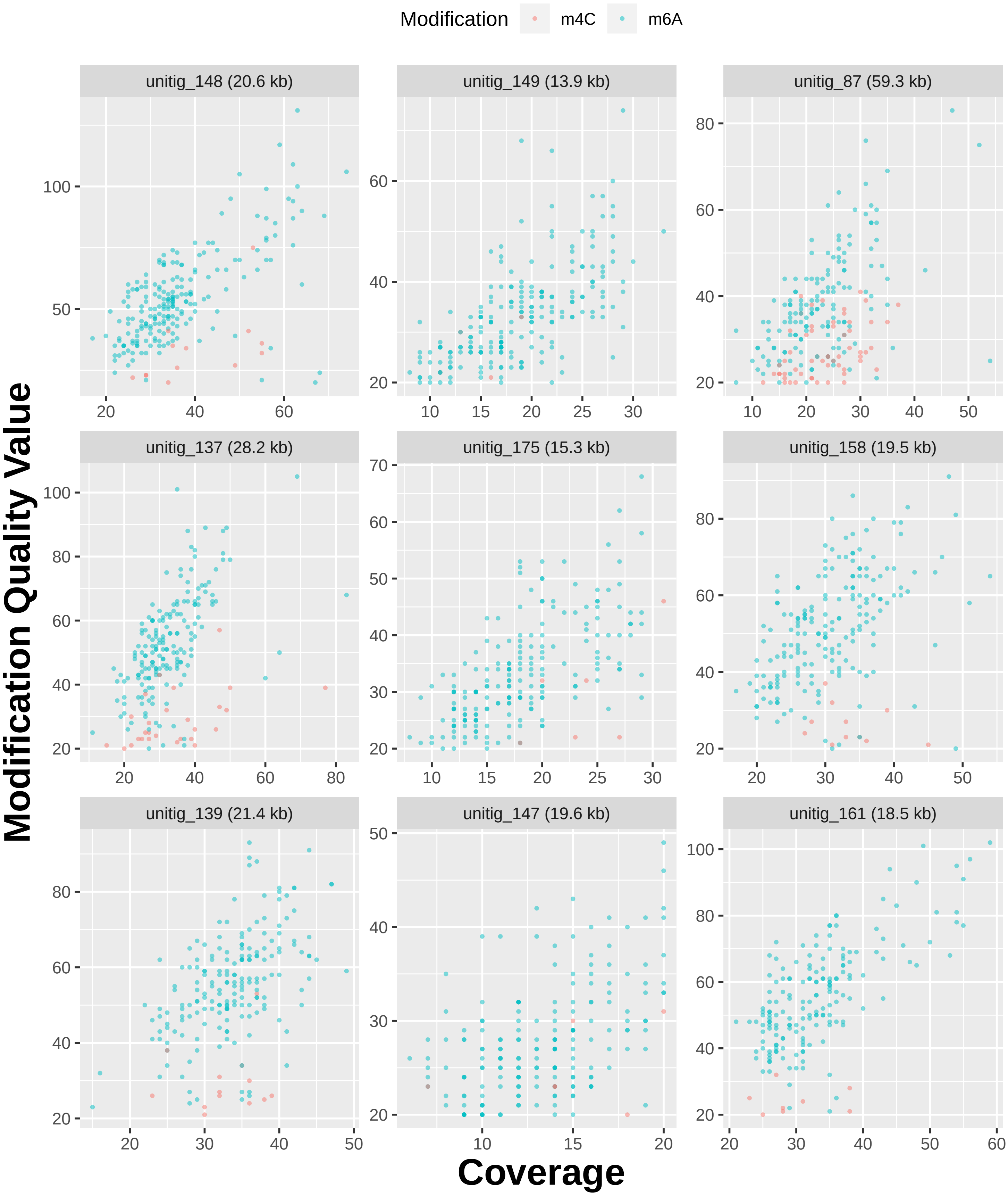

Modification
m4C
m6A
unitig_148 (20.6 kb)
unitig_149 (13.9 kb)
unitig_87 (59.3 kb)
unitig_137 (28.2 kb)
unitig_175 (15.3 kb)
unitig_158 (19.5 kb)
unitig_139 (21.4 kb)
unitig_147 (19.6 kb)
unitig_161 (18.5 kb)
Modification Quality Value
Coverage

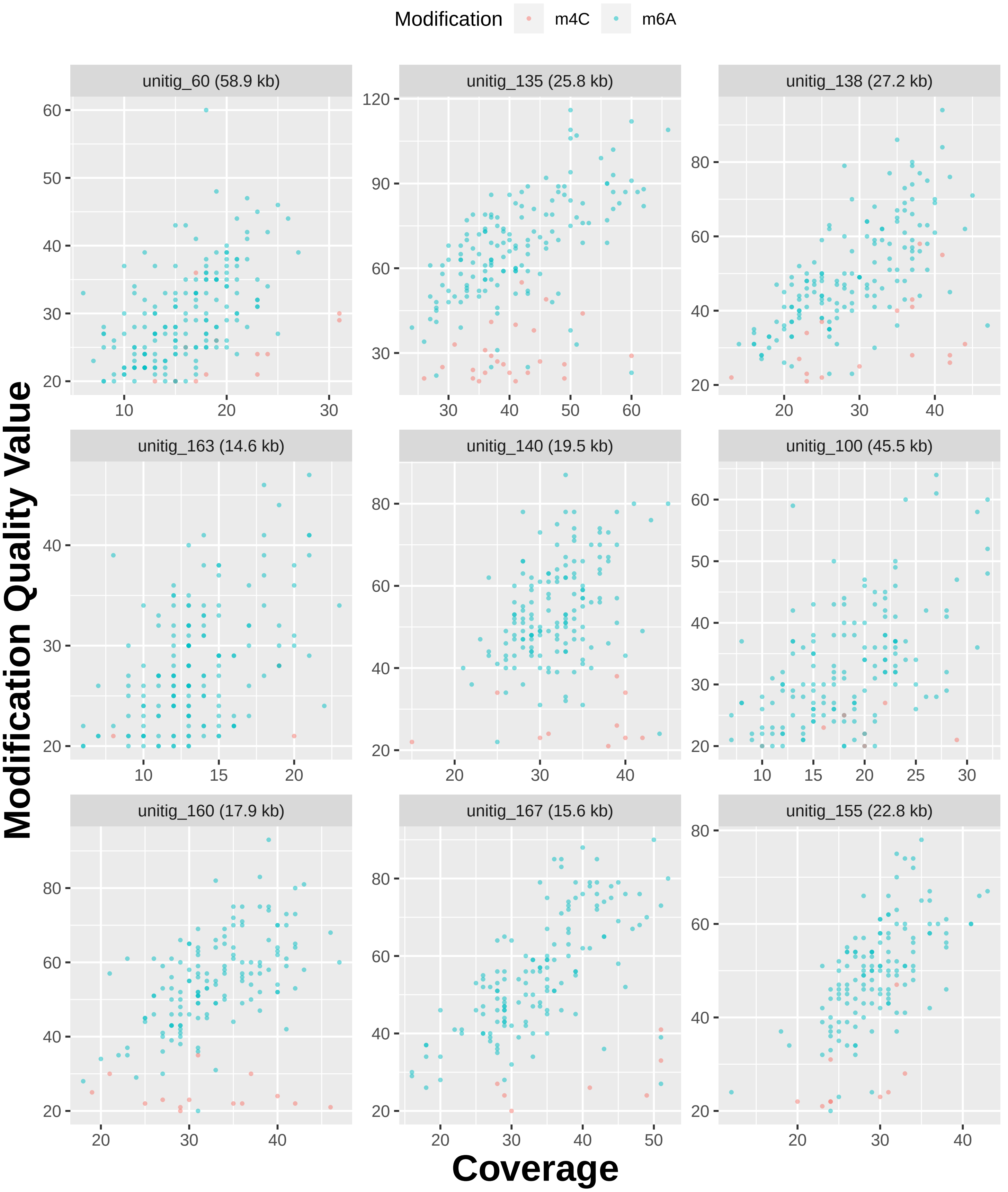

Modification
m4C
m6A
unitig_60 (58.9 kb)
unitig_135 (25.8 kb)
unitig_138 (27.2 kb)
unitig_163 (14.6 kb)
unitig_140 (19.5 kb)
unitig_100 (45.5 kb)
unitig_160 (17.9 kb)
unitig_167 (15.6 kb)
unitig_155 (22.8 kb)
Modification Quality Value
Coverage

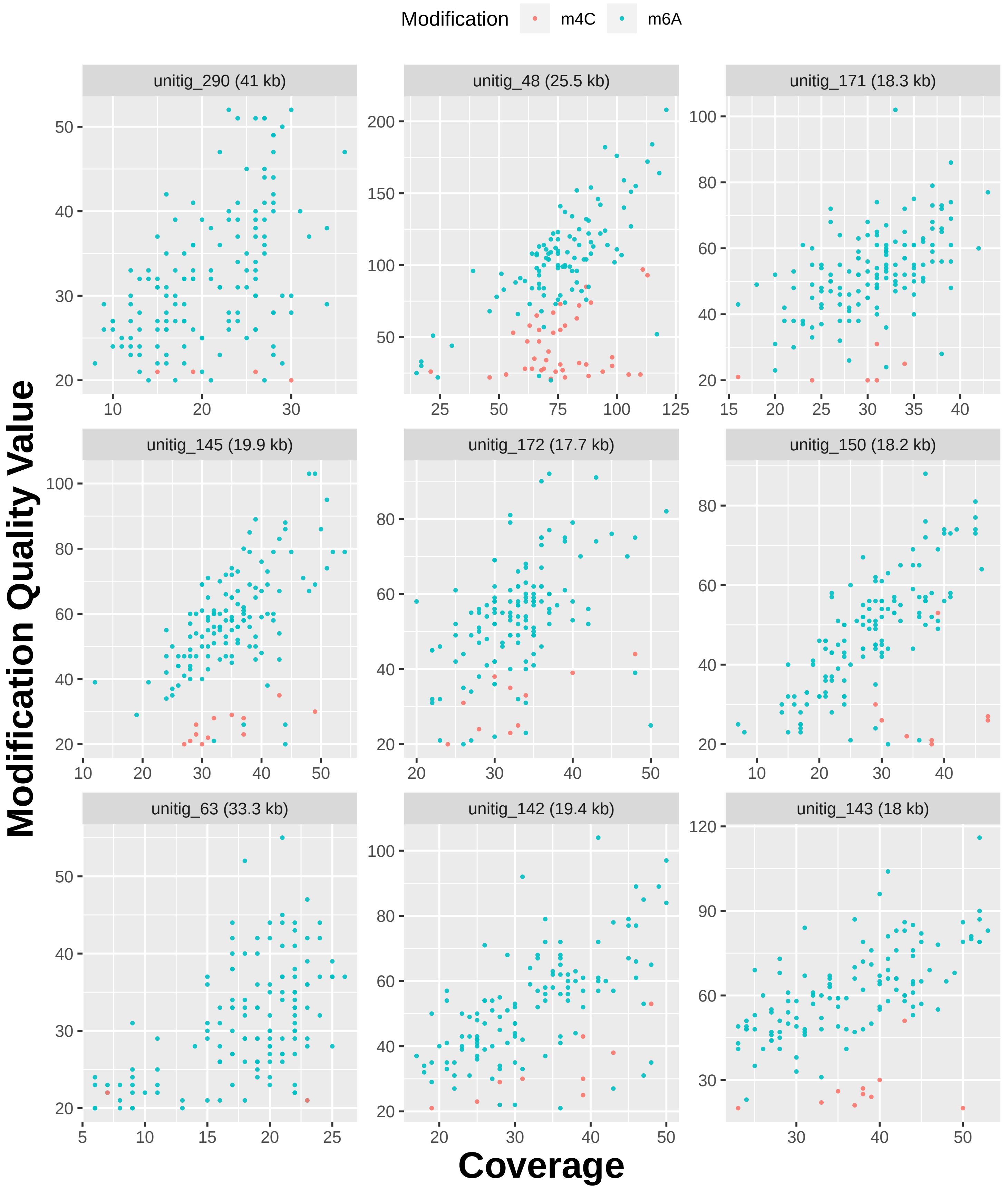

Modification
m4C
m6A
unitig_290 (41 kb)
unitig_48 (25.5 kb)
unitig_171 (18.3 kb)
unitig_145 (19.9 kb)
unitig_172 (17.7 kb)
unitig_150 (18.2 kb)
unitig_63 (33.3 kb)
unitig_142 (19.4 kb)
unitig_143 (18 kb)
Modification Quality Value
Coverage

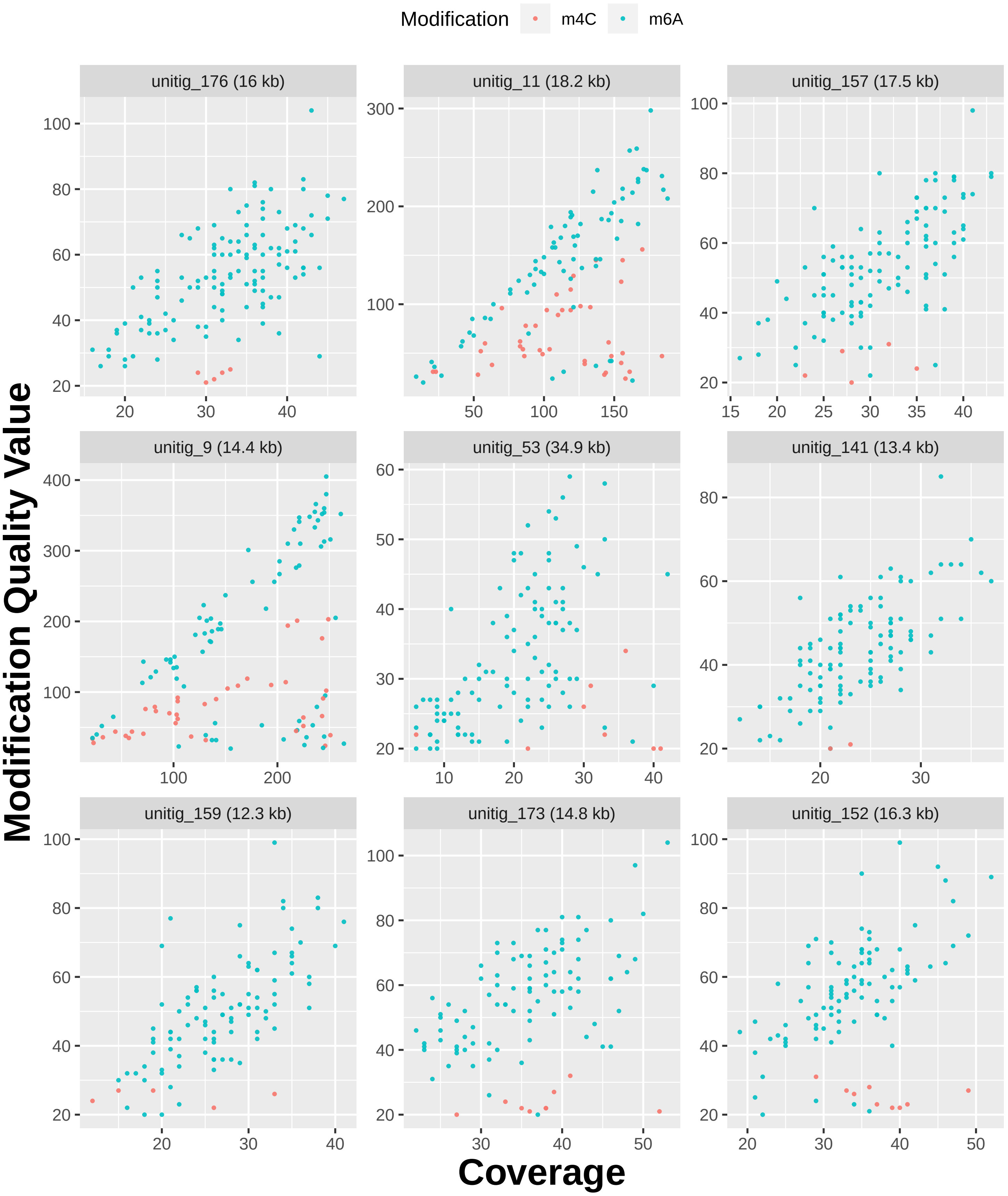

Modification
m4C
m6A
unitig_176 (16 kb)
unitig_11 (18.2 kb)
unitig_157 (17.5 kb)
unitig_9 (14.4 kb)
unitig_53 (34.9 kb)
unitig_141 (13.4 kb)
unitig_159 (12.3 kb)
unitig_173 (14.8 kb)
unitig_152 (16.3 kb)
Modification Quality Value
Coverage

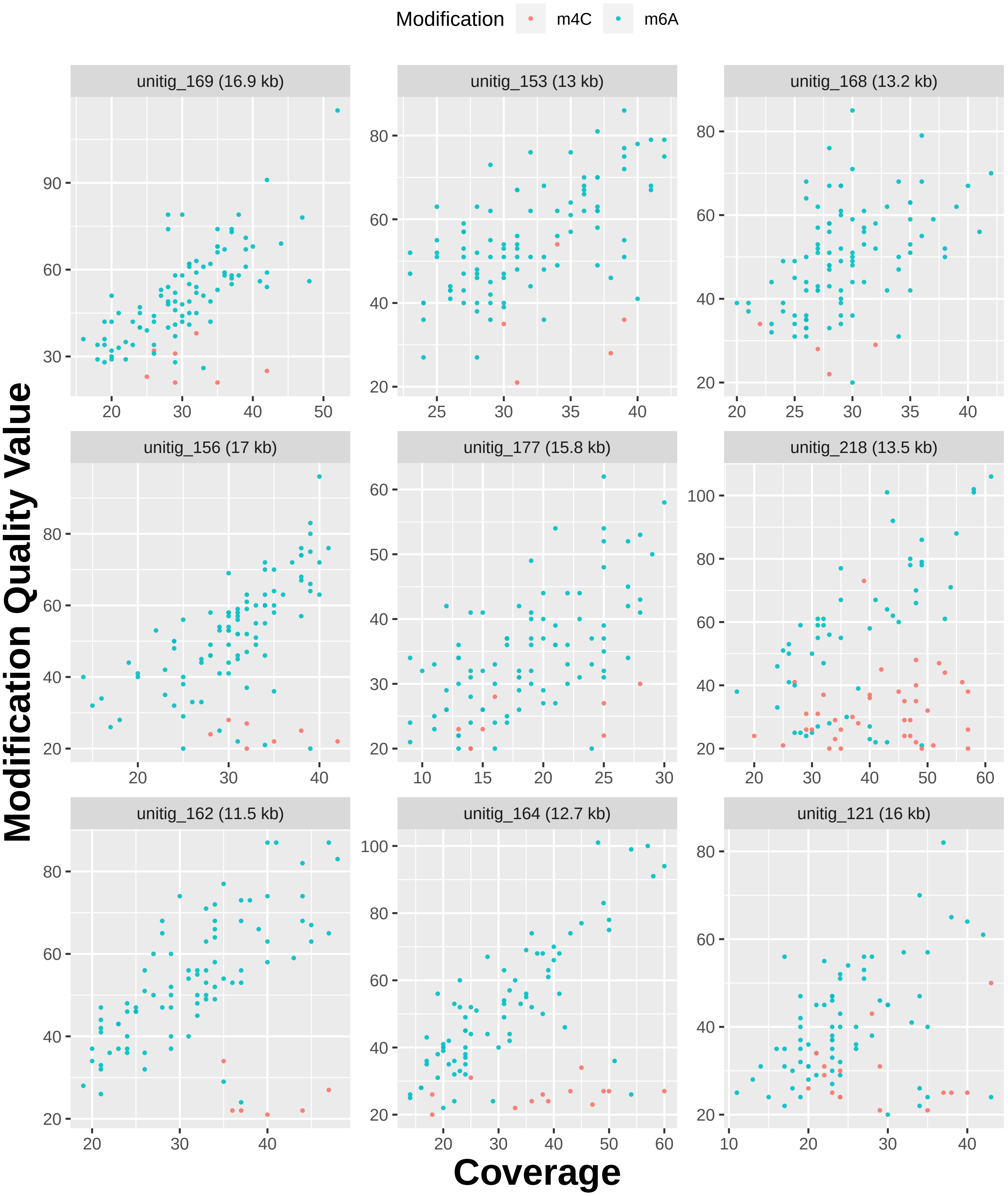

Modification
m4C
m6A
unitig_169 (16.9 kb)
unitig_153 (13 kb)
unitig_168 (13.2 kb)
unitig_156 (17 kb)
unitig_177 (15.8 kb)
unitig_218 (13.5 kb)
unitig_162 (11.5 kb)
unitig_164 (12.7 kb)
unitig_121 (16 kb)
Modification Quality Value
Coverage

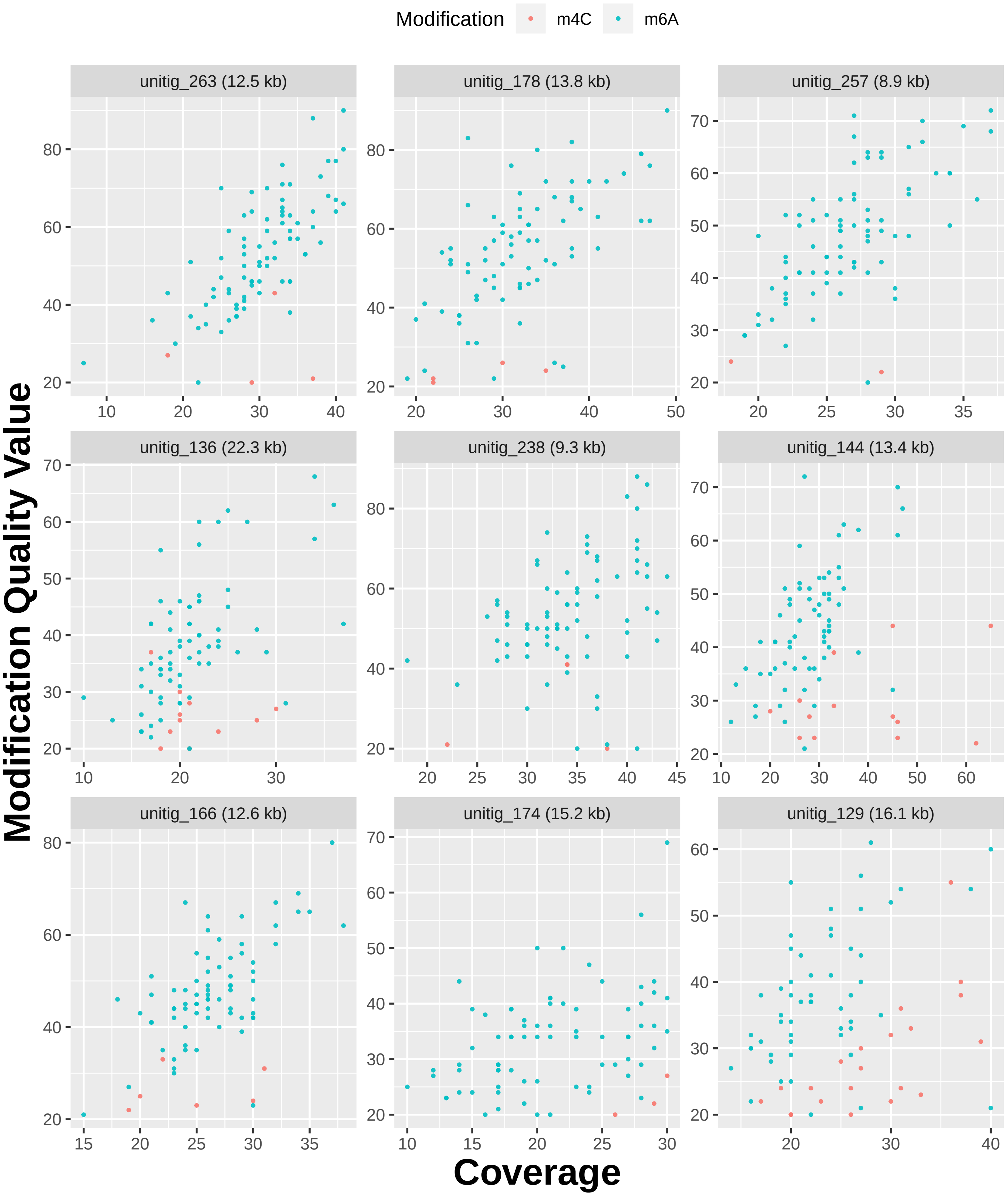
Modification
m4C
m6A
unitig_263 (12.5 kb)
unitig_178 (13.8 kb)
unitig_257 (8.9 kb)
unitig_136 (22.3 kb)
unitig_238 (9.3 kb)
unitig_144 (13.4 kb)
unitig_166 (12.6 kb)
unitig_174 (15.2 kb)
unitig_129 (16.1 kb)
Modification Quality Value
Coverage

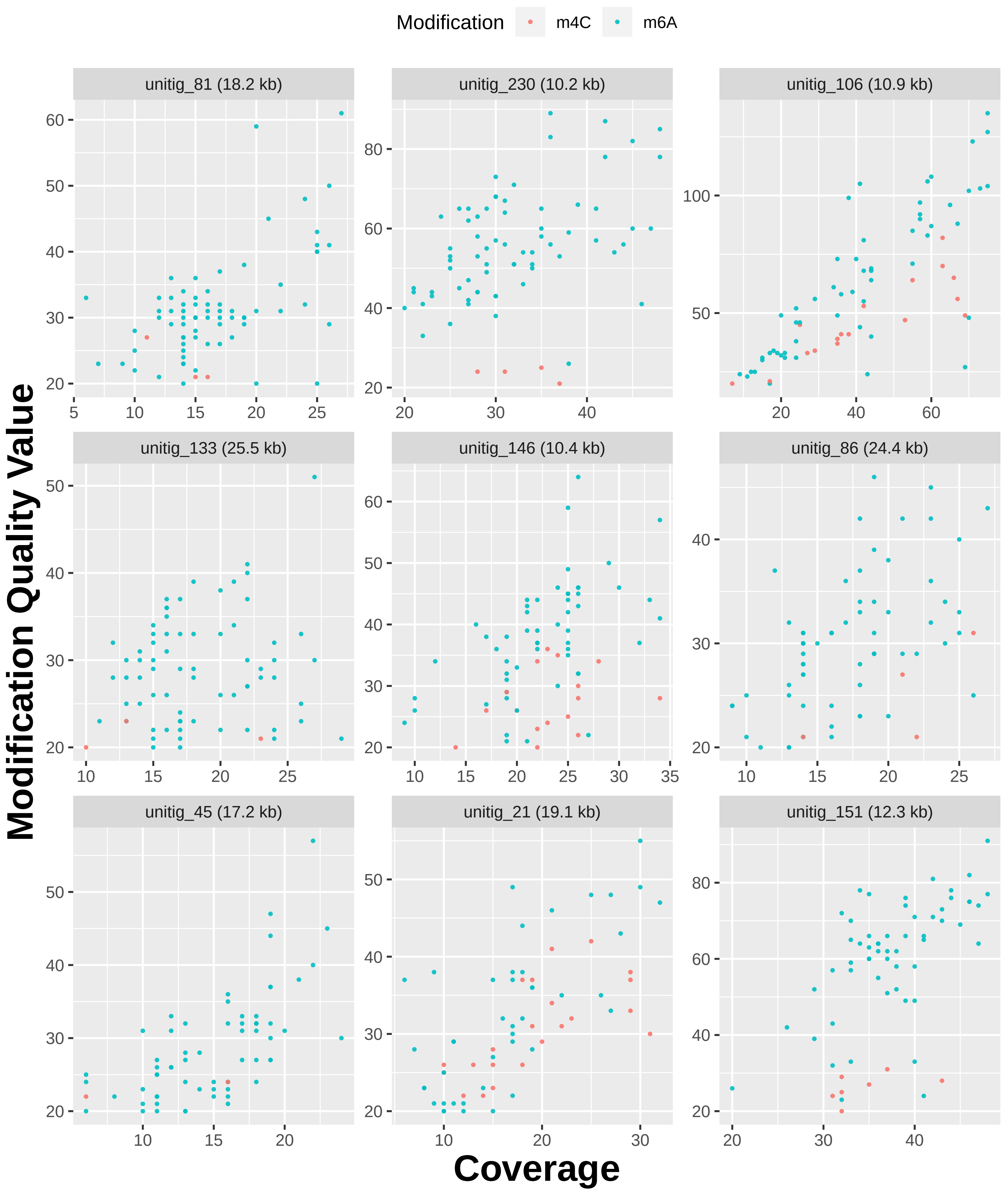
Modification
m4C
m6A
unitig_81 (18.2 kb)
unitig_230 (10.2 kb)
unitig_106 (10.9 kb)
unitig_133 (25.5 kb)
unitig_146 (10.4 kb)
unitig_86 (24.4 kb)
unitig_45 (17.2 kb)
unitig_21 (19.1 kb)
unitig_151 (12.3 kb)
Modification Quality Value
Coverage

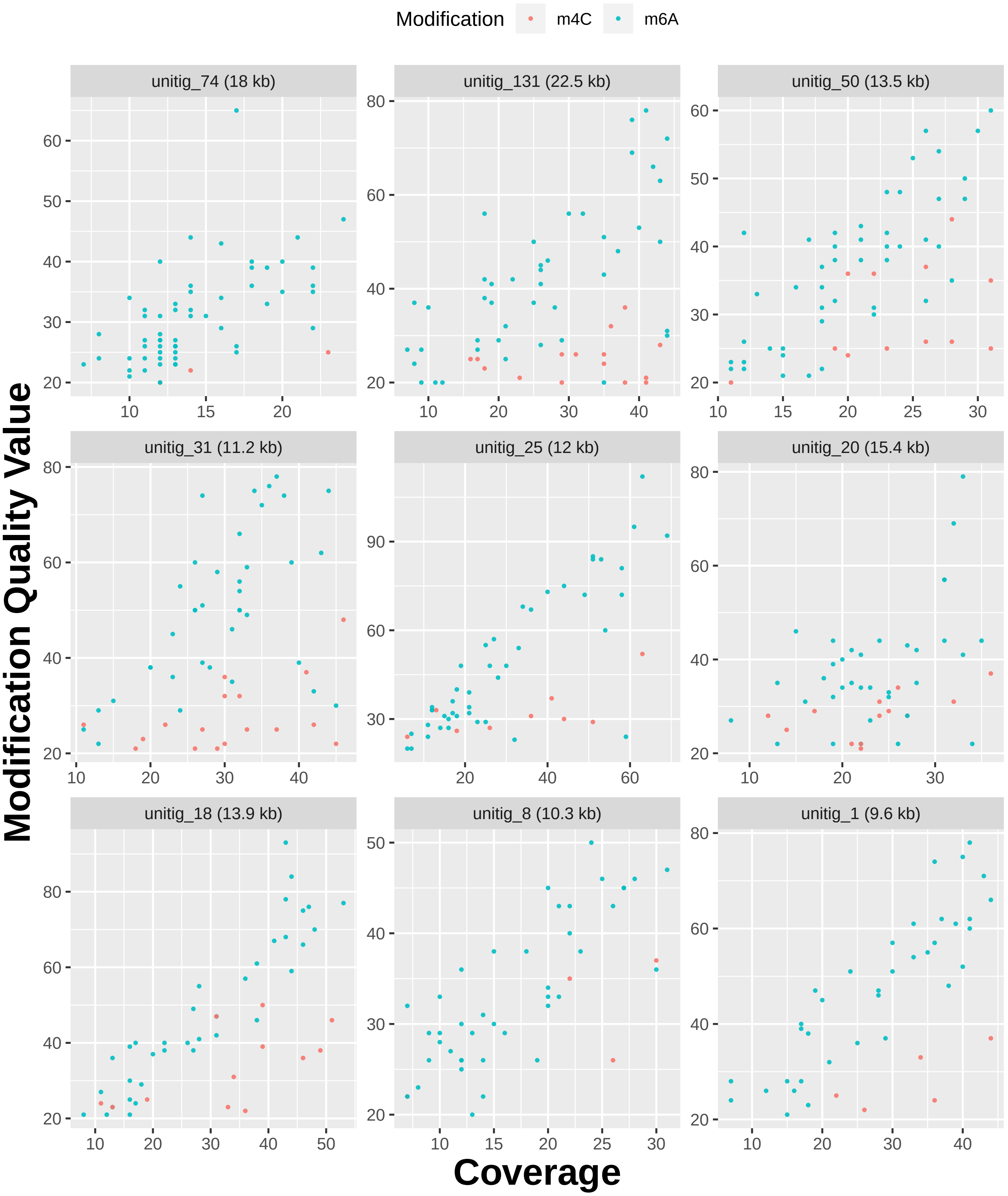
Modification
m4C
m6A
unitig_74 (18 kb)
unitig_131 (22.5 kb)
unitig_50 (13.5 kb)
unitig_31 (11.2 kb)
unitig_25 (12 kb)
unitig_20 (15.4 kb)
unitig_18 (13.9 kb)
unitig_8 (10.3 kb)
unitig_1 (9.6 kb)
Modification Quality Value
Coverage

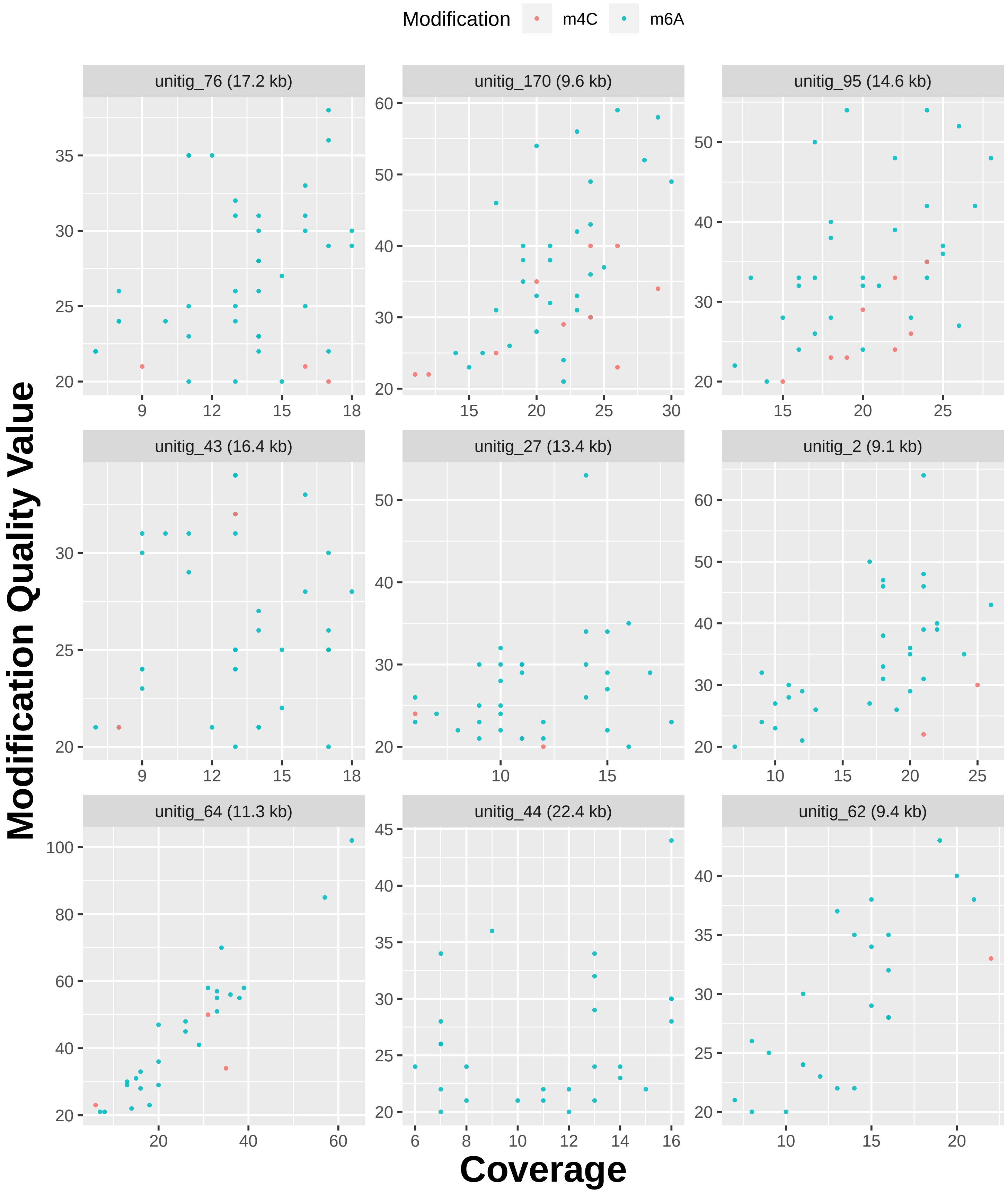

Modification
m4C
m6A
unitig_76 (17.2 kb)
unitig_170 (9.6 kb)
unitig_95 (14.6 kb)
unitig_43 (16.4 kb)
unitig_27 (13.4 kb)
unitig_2 (9.1 kb)
unitig_64 (11.3 kb)
unitig_44 (22.4 kb)
unitig_62 (9.4 kb)
Modification Quality Value
Coverage

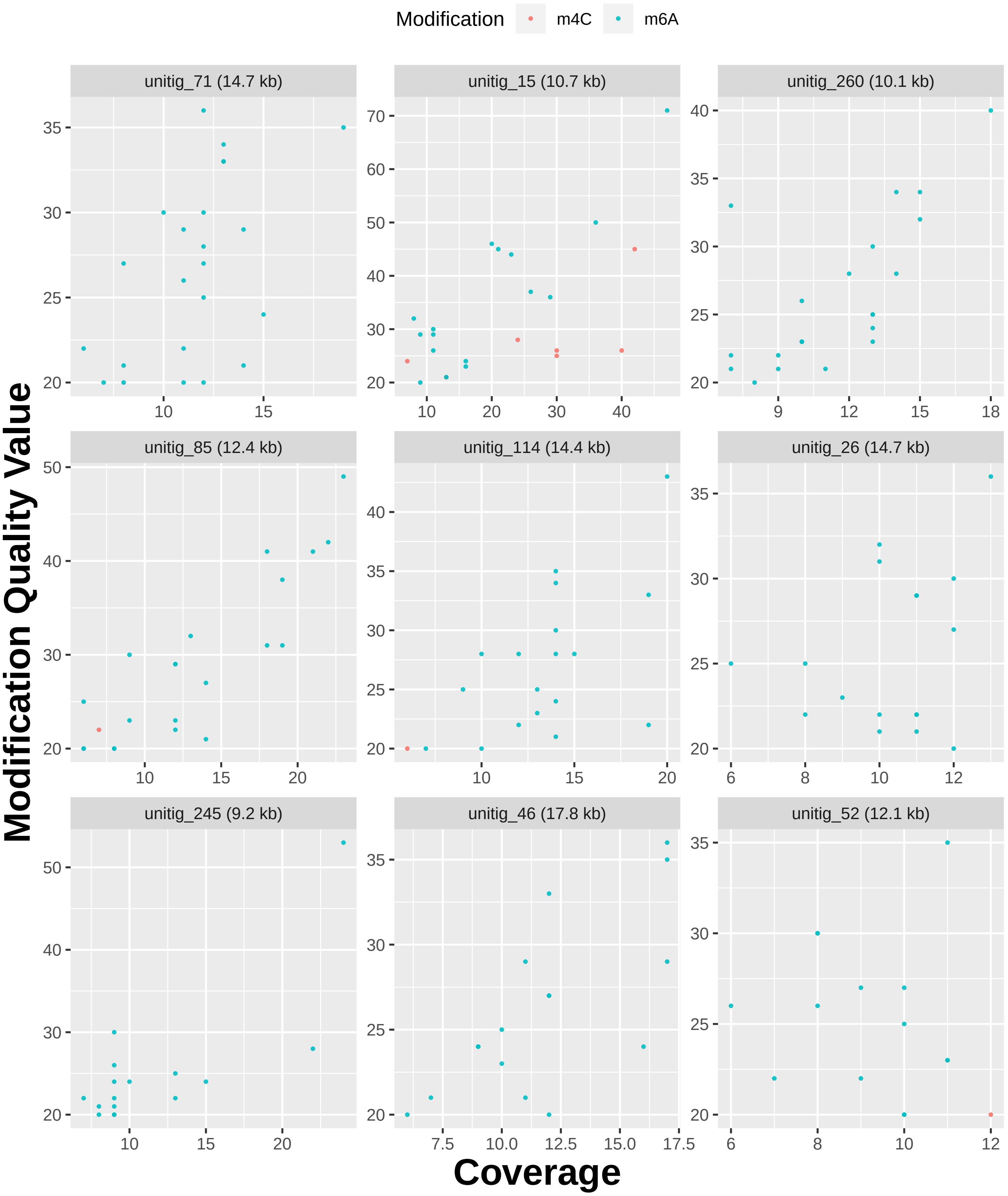
Modification
m4C
m6A
unitig_71 (14.7 kb)
unitig_15 (10.7 kb)
unitig_260 (10.1 kb)
unitig_85 (12.4 kb)
unitig_114 (14.4 kb)
unitig_26 (14.7 kb)
unitig_245 (9.2 kb)
unitig_46 (17.8 kb)
unitig_52 (12.1 kb)
Modification Quality Value
Coverage

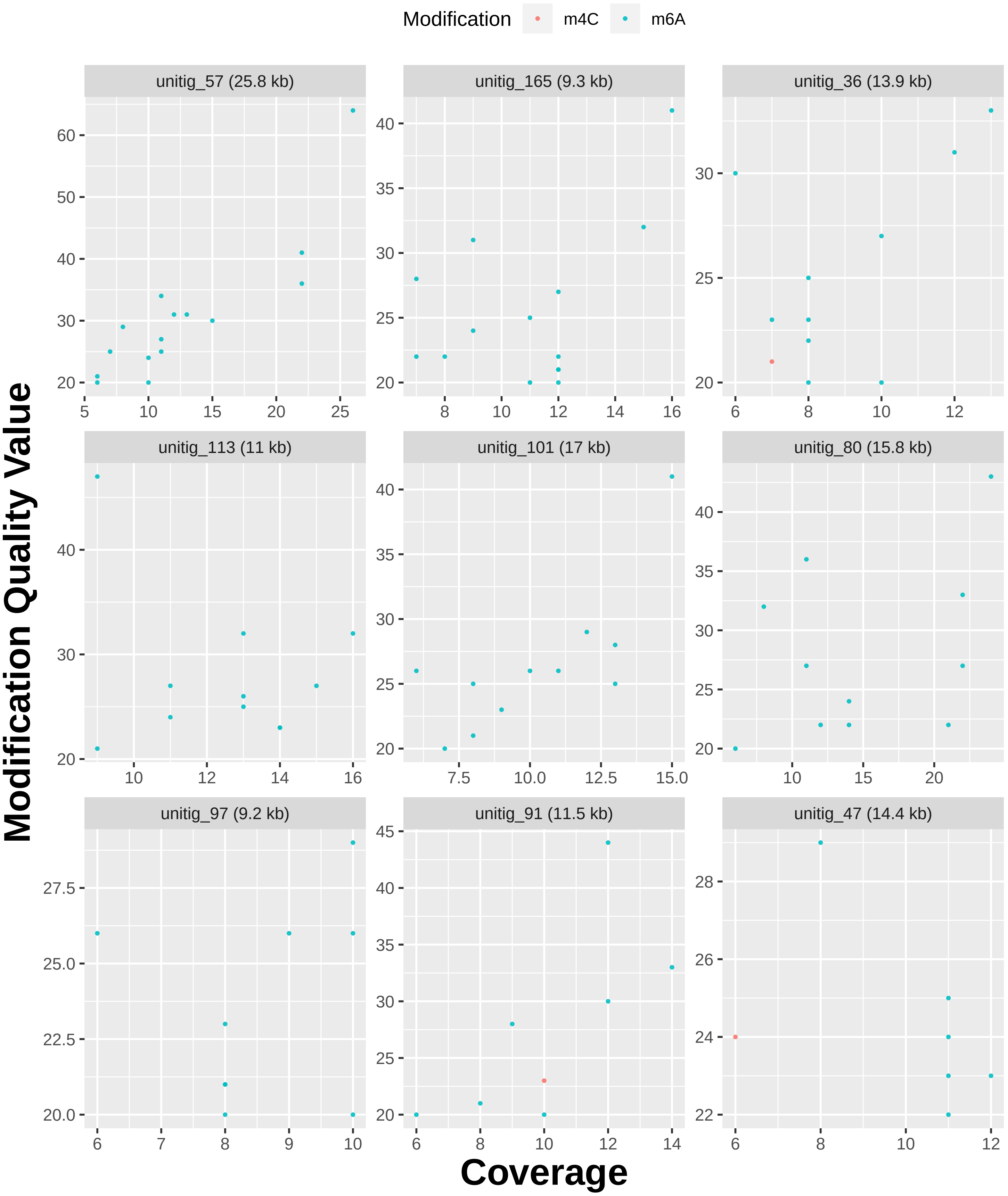
Modification
m4C
m6A
unitig_57 (25.8 kb)
unitig_165 (9.3 kb)
unitig_36 (13.9 kb)
unitig_113 (11 kb)
unitig_101 (17 kb)
unitig_80 (15.8 kb)
unitig_97 (9.2 kb)
unitig_91 (11.5 kb)
unitig_47 (14.4 kb)
Modification Quality Value
Coverage

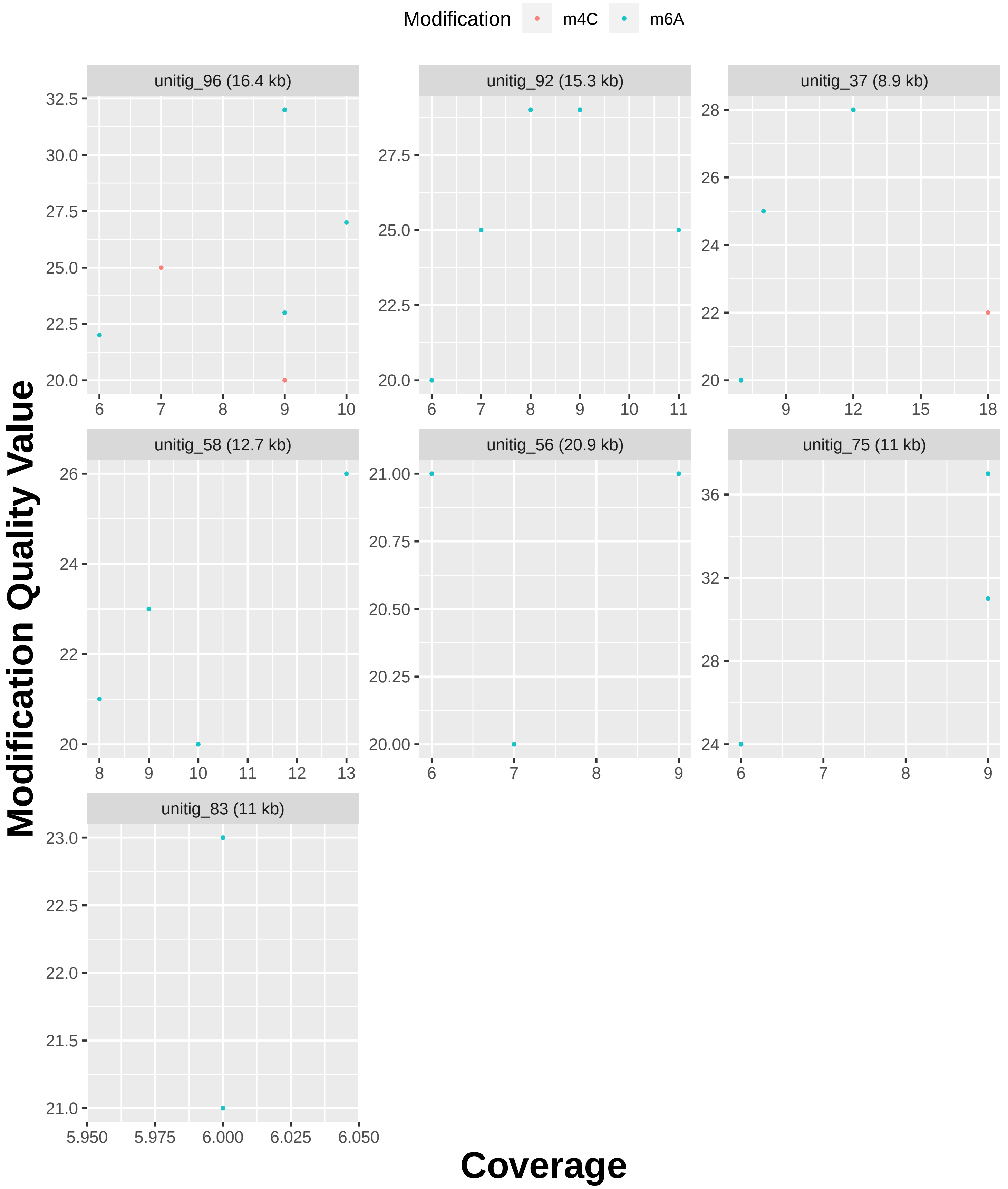

Modification
m4C
m6A
unitig_96 (16.4 kb)
unitig_92 (15.3 kb)
unitig_37 (8.9 kb)
unitig_58 (12.7 kb)
unitig_56 (20.9 kb)
unitig_75 (11 kb)
unitig_83 (11 kb)
Modification Quality Value
Coverage
